# T2R bitter taste receptors regulate apoptosis and may be associated with survival in head and neck squamous cell carcinoma

**DOI:** 10.1101/2021.05.17.444527

**Authors:** Ryan M. Carey, Derek B. McMahon, Karthik Rajasekaran, Indiwari Gopallawa, Jason G. Newman, Devraj Basu, Elizabeth A. White, Robert J. Lee

**Affiliations:** Department of Otorhinolaryngology, University of Pennsylvania Perelman School of Medicine, Philadelphia, Pennsylvania, USA; Department of Physiology, University of Pennsylvania Perelman School of Medicine, Philadelphia, Pennsylvania, USA

**Author notes:** To whom correspondence should be addressed: Ryan M. Carey or Robert J. Lee, Department of Otorhinolaryngology—Head and Neck Surgery, Hospital of the University of Pennsylvania, 3400 Spruce Street, 5^th^ floor Ravdin Suite A, Philadelphia, PA 19104; (R.M.C.) or, (R.J.L.).

**Keywords:** G protein-coupled receptor (GPCR), taste receptors, head and neck squamous cell carcinoma, apoptosis, calcium, cell signaling, genetics

## Abstract

Better management of head and neck squamous cell carcinomas (HNSCCs) requires a clearer understanding of tumor biology and disease risk. Bitter taste receptors (T2Rs) have been studied in several cancers, including thyroid, salivary, and GI, but their role in HNSCC has not been explored. We found that HNSCC patient samples and cell lines expressed functional T2Rs on both the cell and nuclear membranes. Bitter compounds, including bacterial metabolites, activated T2R-mediated nuclear Ca^2+^ responses leading to mitochondrial depolarization, caspase activation and ultimately apoptosis. Buffering nuclear Ca^2+^ elevation blocked caspase activation. Furthermore, increased expression of T2Rs in HNSCCs from The Cancer Genome Atlas (TCGA) is associated with improved overall survival. This work suggests that T2Rs are potential biomarkers to predict outcomes and guide treatment selection, may be leveraged as therapeutic targets to stimulate tumor apoptosis, and may mediate tumor-microbiome crosstalk in HNSCC.

## Introduction

Head and neck squamous cell carcinoma (HNSCC) is the 6th most common cancer worldwide, with an expected 30% incidence increase by 2030 [1]. HNSCC often presents with locally advanced disease [2], and approximately half of patients die ≤5 years after diagnosis [3]. Treatment is based on clinical and pathologic risk factors, typically consisting of a combination of surgery, radiation, chemotherapy, and immunotherapy in select cases [4, 5]. Because treatment side-effects impact quality of life, it is important to tailor the aggressiveness of therapy to disease risk [6].

HNSCCs associated with human papillomavirus (HPV) show improved overall survival and recurrence-free survival compared to HPV-negative tumors. Thus, HPV status is an important tool for risk stratification and treatment selection [7, 8]. There is need for additional methods or biomarkers for stratifying disease risk. There is also a need for alternative therapies that maximize survival while minimizing morbidity in HNSCC. The ideal cancer therapy may be one that exploits cellular machinery to prevent immune system evasion or impact tumor microenvironment.

The current study suggests that bitter taste receptors (T2Rs) may be important for risk stratification and/or as novel therapeutic targets in HNSCC. Initially identified on the tongue, T2Rs are expressed in other tissues, including the gastrointestinal (GI) [9] and airway epithelia [10, 11]. There are 25 human T2R isoforms [11] encoded by *TAS2R* genes [12]. T2Rs are G protein-coupled receptors (GPCRs) that signal through Gα-mediated cAMP decrease and Gβγ-activation of phospholipase C (PLC) and calcium (Ca^2+^) release [11]. T2Rs serve a diverse array of chemosensory functions, including in innate immunity [11, 13]. For example, T2Rs in nasal cells bind bacterial products to activate Ca^2+^-driven nitric oxide (NO) production to increase cilia beating and kill bacteria [10, 11].

T2Rs have also been explored in some cancers [14-23], including thyroid [22], salivary [23], and GI cancers [14-19]. Polymorphisms in the *TA2R38* gene encoding the T2R38 receptor are associated with elevated cancer risk [16-18, 24]. Individuals with specific *TAS2R3* and *4* haplotypes may have lower risk of papillary thyroid cancer [22]. Recent work demonstrated elevated expression of many GPCRs in solid tumors compared with normal tissue [25]. While there is prior work on T2Rs in cancer, an association between T2Rs and HNSCC and the potential T2R signaling pathways within cancer cells are unknown.

We hypothesized that HNSCCs may differentially express *TAS2Rs* compared to normal tissue and some T2Rs are functional and regulate cellular processes. We characterized *TAS2R* expression in HNSCC patient samples and cell lines. We demonstrate that some T2Rs have intracellular or intranuclear localization and can be activated to cause mitochondrial depolarization and apoptosis in HNSCC cells. We subsequently evaluated HNSCC patients for association of *TAS2R* expression levels with survival outcomes using The Cancer Genome Atlas (TCGA).

## Materials and Methods

Unless noted, all reagents and protocols were used as previously described [10, 26-29]. All reagents and catalogue numbers are in **Supplementary Table S1**.

### Cell culture

SCC4, SCC15, SCC90, and SCC152 cells were from ATCC (Manassas, VA USA). UMSCC47 (SCC47) was from Sigma-Aldrich (St. Louis, MO USA). OCTT2 cell line was derived from a surgical specimen of an oral SCC tumor [30]. VU147T was from Dr. Hans Joenje, VU Medical Center, Netherlands. All cancer cell lines were grown in submersion in high glucose Dulbecco’s Modified Eagle Medium (Gibco; Gaithersburg, MD USA) plus 10% FBS, penicillin/streptomycin mix (Gibco), and non-essential amino acids (Gibco). Primary gingival keratinocytes were from ATCC (Manassas, VA USA) and used within 2 passages using dermal cell basal keratinocyte medium (ATCC).

### Patient samples

SCC specimens were obtained from patients undergoing diagnostic biopsies of HNSCC tumors as part of routine clinical care (University of Pennsylvania Institutional Review Board protocol #417200). Tissue acquisition was done in accordance with the University of Pennsylvania guidelines for the use of residual clinical material and in accordance with the U.S. Department of Health and Human Services code of federal regulation Title 45 CFR 46.116 and the Declaration of Helsinki. Tumor and contralateral normal tissue was obtained. Specimens were divided for pathologic evaluation and expression analysis samples were collected in TRIzol (ThermoFisher Scientific).

All tumor specimens had final pathology consistent with SCC. HPV-positivity was determined after p16 testing, based-on immunohistochemistry that demonstrated ≥70% of tumor nuclear and cytoplasmic staining [31]. Testing was performed on cancers of the oropharynx, but not for cancers of other HNSCC sites per the University of Pennsylvania guidelines.

### Quantitative reverse-transcription PCR

Patient samples and subconfluent cultures were resuspended in TRIzol (ThermoFisher Scientific). RNA was isolated and purified (Direct-zol RNA kit; Zymo Research), reverese transcribed via High-Capacity cDNA Reverse Transcription Kit (ThermoFisher Scientific), and quantified using TaqMan qPCR probes for the 25 *TAS2R* genes and UBC1 (QuantStudio 5; ThermoFisher Scientific).

### Live cell imaging

Cells were loaded with 5 μM of Fluo-4-AM or Fluo-8-AM for 45 minutes at room temperature in the dark, then imaged using an Olympus IX-83 microscope (20x 0.75 NA PlanApo objective), FITC fitlers, Orca Flash 4.0 sCMOS camera (Hamamatsu, Tokyo, Japan), MetaFluor (Molecular Devices, Sunnyvale, CA USA), and XCite 120 LED Boost (Excelitas Technologies). For nuclear Ca^2+^, R-GECO-nls [32] was transfected using Lipofectamine 3000 (ThermoFisher Scientific) 24-48 hours prior to imaging. Live cell images were taken as above with standard TRITC filters.

For Flip-GFP, cells were transfected and imaging was carried out on an Olympus IX-83 microscope as above with 10x (0.4 NA) or 4x (0.16 NA) objective and FITC and TRITC filter sets. For CFP-DEVD-mVenus, cells were imaged at 40x (0.75NA objective) with CFP/YFP filters (Chroma 89002-ET-ECFP/EYFP) in excitation and emission filter wheels (Sutter Lambda LS). Images were acquired at 37°C using a stage-top incubator (Tokai Hit, Tokyo Japan).

### Immunofluorescence

Cultures were fixed in 4% paraformaldehyde for 20 min at room temperature, followed by blocking and permeabilization in DPBS containing 5% normal donkey serum, 1% BSA, 0.2% saponin, and 0.1% Triton X-100 for 45 min. Cultures were with T2R or tubulin antibodies (1:100) at 4°C overnight. Several T2R antibodies (T2R14, T2R4) were validated previously [10, 13, 33]. Cultures were then incubated with AlexaFluor-labeled donkey secondary antibodies (1:1000) at 4°C for 1 hour, then mounted with Fluoroshield with DAPI (Abcam). Images were obtained using an Olympus IX-83 microscope (60x 1.4 NA oil; MetaMorph software).

### Mitochondrial membrane potential and apoptosis measurements

JC-1 dye was added to sub-confluent cells on 24-well glass bottom-plates (CellVis) 10 min prior to measurements (ex.488/em.535 and em.590). CellEvent Caspase 3/7 was added directly prior to measurements (ex.495/em.540) per manufacturer’s specifications. XTT was added directly prior to measurements at 475nm and 660nm. All data for JC-1, CellEvent Caspase 3/7, and XTT assays were obtained Tecan (Männedorf, Switzerland) Spark 10M. TMRE, Red Dot 2, and Hoechst were added per manufacturer’s specifications. Data were obtained using an Olympus IX-83 microscope (20x 0.8 NA objective; MetaFluor) with DAPI, TRITC, and Cy5 filters.

### The Cancer Genome Atlas analysis (TCGA)

Data was obtained from Firehose Legacy, The Cancer Genome Atlas (TCGA) from cBio Cancer Genomics Portal (cbioportal.org) [34, 35]. This database compares each queried gene to that gene’s expression in a reference population, which typically consists of all samples that are diploid for the gene [35]. We included all HNSCC samples with mutation and copy number alteration (CNA) data available. The database was queried for all 25 *TAS2R* genes. Analysis used cBio Portal software.

### Data analysis and statistics

T-tests (two comparisons only) and one-way ANOVA (>2 comparisons) were calculated using GraphPad Prism with appropriate post-tests, as indicated. Additional data analysis was performed using Microsoft Excel. All figures used the following annotations: *p* < 0.05 (*), *p* < 0.01 (**), *p* < 0.001 (***), and no statistical significance (ns). All data points represent the mean ± standard error of the mean.

## RESULTS

### T2Rs are differentially expressed and localized in HNSCC

We examined expression of 25 *TAS2Rs* in HNSCC tissue from patients undergoing diagnostic biopsies during routine clinical care. Tissue was obtained from the tumor site and corresponding contralateral normal site from 10 patients (**Supplementary Table S2**). Quantitative reverse transcription PCR (qPCR) demonstrated variable expression of *TAS2R*s (**Fig. 1A-B**). In aggregate, there were no significant differences in expression of individual *TAS2R*s between control and cancer samples (**Fig. 1A**).

**Fig. 1.**
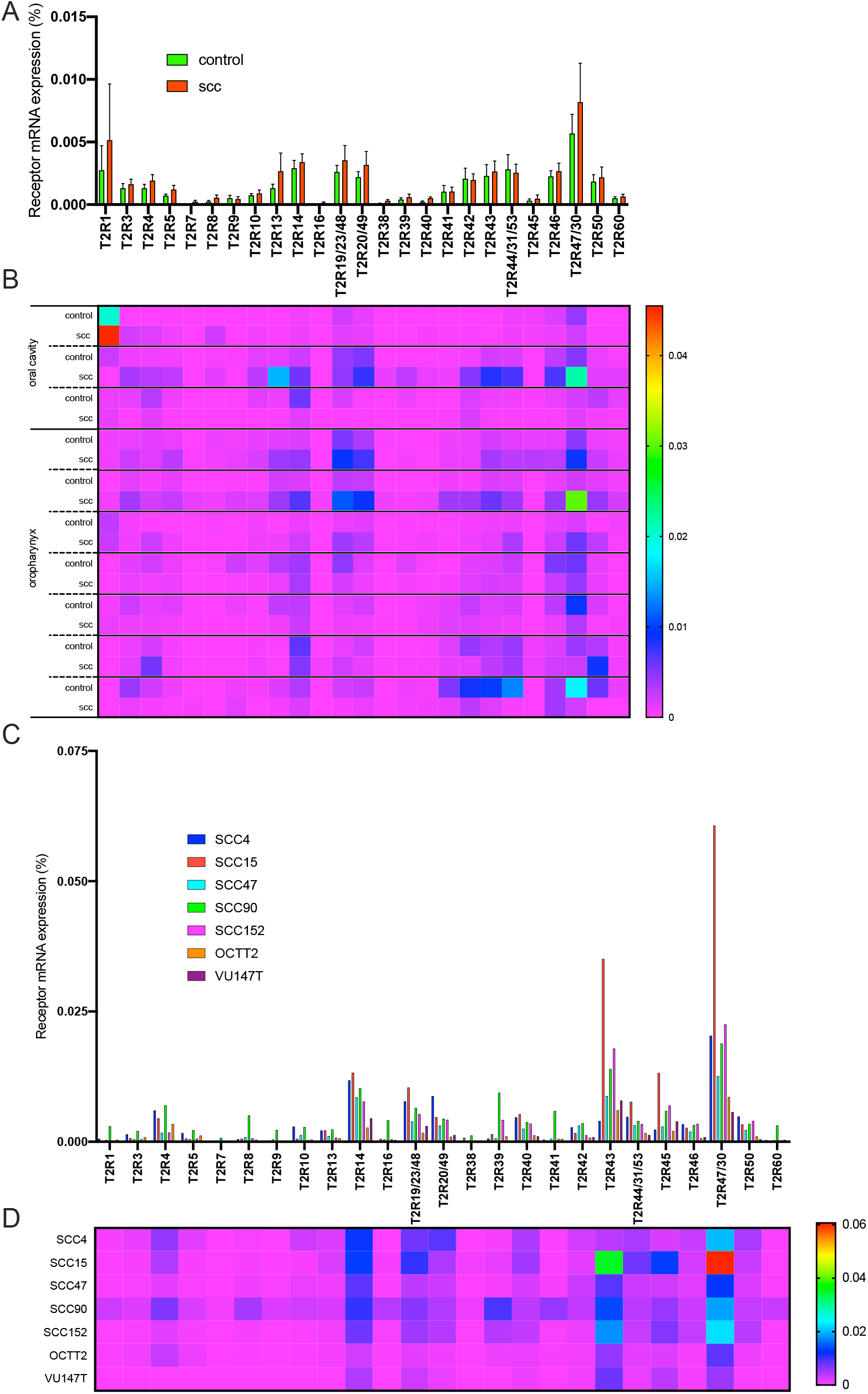
There is variable mRNA expression of bitter (T2R) taste receptor genes in head and neck squamous cell carcinoma (HNSCC). **A** Quantitative PCR (qPCR) expression analysis of T2R transcripts from biopsy specimens of HNSCC (“cancer”) compared to contralateral normal tissue (“control”) from the same patient (mean ± SEM; 10 patients). Relative expression is normalized to ubiquitin C (UBC) housekeeping gene. No significant differences in individual T2R gene expression between cancer and control samples by 2-way ANOVA with Bonferroni posttest. **B** Heatmap representation of the same T2R transcript data, grouped by individual patients. **C** Bar graph and **D** heatmap of qPCR T2R transcripts in HNSCC cell lines SCC4, SCC15, SCC47, SCC90, SCC152, OCTT2, and VU147T. Relative expression is normalized to UBC.

However, comparison at the individual patient level demonstrated that some individuals had increased *TAS2R* expression in the tumor specimens while others had decreased expression compared to matched control tissue (**Fig. 1B**). This small sample size suggests that T2Rs are expressed in HNSCC tissue, and that further follow-up study with larger patient populations is warranted to determine differences in normal vs cancer tissue, including potential associations with clinical outcomes.

*TAS2R* expression was measured in HNSCC cell lines SCC4, SCC15, SCC47, SCC90, SCC152, OCTT2, and VU147T (**Fig. 1C, D**). Like patient samples, cells had variable *TAS2R* expression, but specific *TAS2R*s were consistently high, including *TAS2R4, TAS2R14, TAS2R19* (also known as *TAS2R23* or *TAS2R48*), *TAS2R20* (also known as *TAS2R49*), and *TAS2R30* (formerly known as *TAS2R47*), *TAS2R43* and *TAS2R45*. Similarities in *TAS2R* expression between cell lines and tumor specimens suggest these cells may be useful for studying T2R signaling in HNSCC.

Next, we used confocal immunofluorescence microscopy to visualize T2R localization. Fixed SCC47 and SCC4 cells were stained with antibodies targeting endogenous tubulin and T2Rs (**Fig. 2, Supplementary Fig. S1**). Interestingly, T2R42 localized to the nucleus in both cell lines and T2R13 was nuclear-localized in SCC47 but not SCC4. Other T2Rs (including 4, 8, 10, 14, 30/47, and 46) appeared to localize to intracellular membranes, potentially including the endoplasmic reticulum. T2R expression on the nuclear membrane is novel, and fits with our studies of airway cells, where squamous de-differentiation promotes intracellular and even nuclear localization (preprint [36]).

**Fig. 2.**
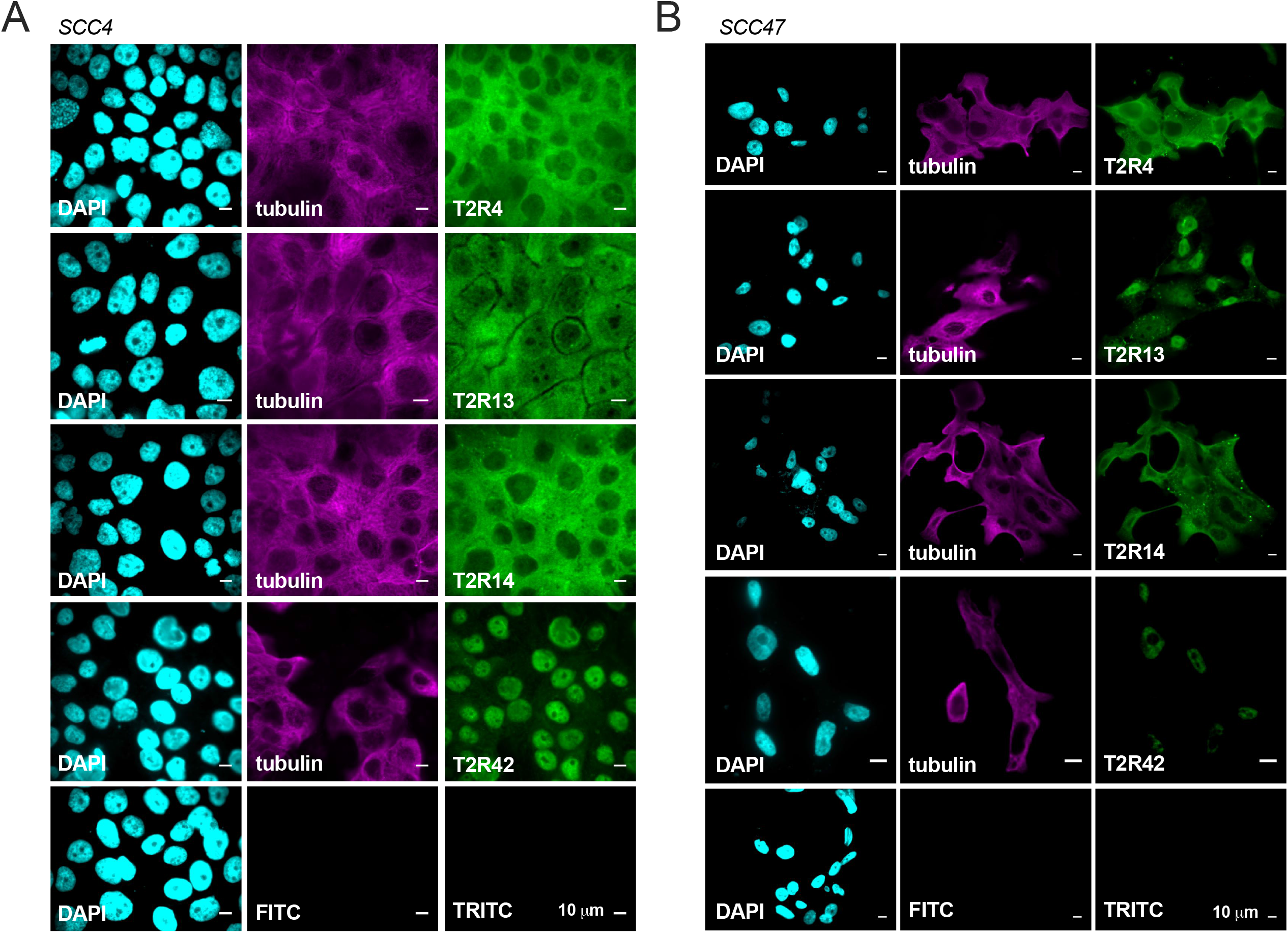
A subset of T2Rs localize to the nucleus of two head and neck squamous cell carcinoma (HNSCC) cell lines. Fixed cultures of HNSCC cell lines **A** SCC4 and **B** SCC47 stained with antibodies targeting endogenous proteins demonstrate nuclear localization of T2R42. T2R13 appears to localize strongly to the nucleus in SCC47 but not in SCC4. T2Rs 4 and 14 are expressed in both cell lines and appear to localize to the plasma membrane. For all images, 1 representative image from 3 experiments were shown. Each antibody was compared to secondary only control at the same microscope settings.

### T2R-activated nuclear Ca^2+^ responses in HNSCC

To determine if T2Rs in HNSCC cells are functional, we examined agonist-induced intracellular calcium (Ca^2+^_i_) changes in SCC47, SCC4, SCC15, SCC152, SCC90, OCT22, and VU147T loaded with Ca^2+^ indicator Fluo-4 (**Fig. 3A**). We tested bitter compounds targeting multiple T2Rs (**Fig. 3B**). SCC4 and SCC47 exhibited Ca^2+^_i_ elevations in response to bitter agonists denatonium benzoate, quinine, diphenidol, flufenamic acid, parthenolide, thujone, and the *P. aeruginosa* quorum-sensing molecule N-3-oxo-dodecanoyl-L-homoserine lactone (3-oxo-C12HSL, 100 μM) (**Fig. 3A-D, Supplementary Fig. S2**). Similarly, SCC15, SCC152, SCC90, OCT22, and VU147T exhibited Ca^2+^_i_ responses to a smaller subset of screened bitter agonists (**Supplementary Fig. S3**). None of the cells responded to T2R38 agonist phenylthiocarbamide (PTC), consistent with *TAS2R38* expression at very low levels.

**Fig. 3.**
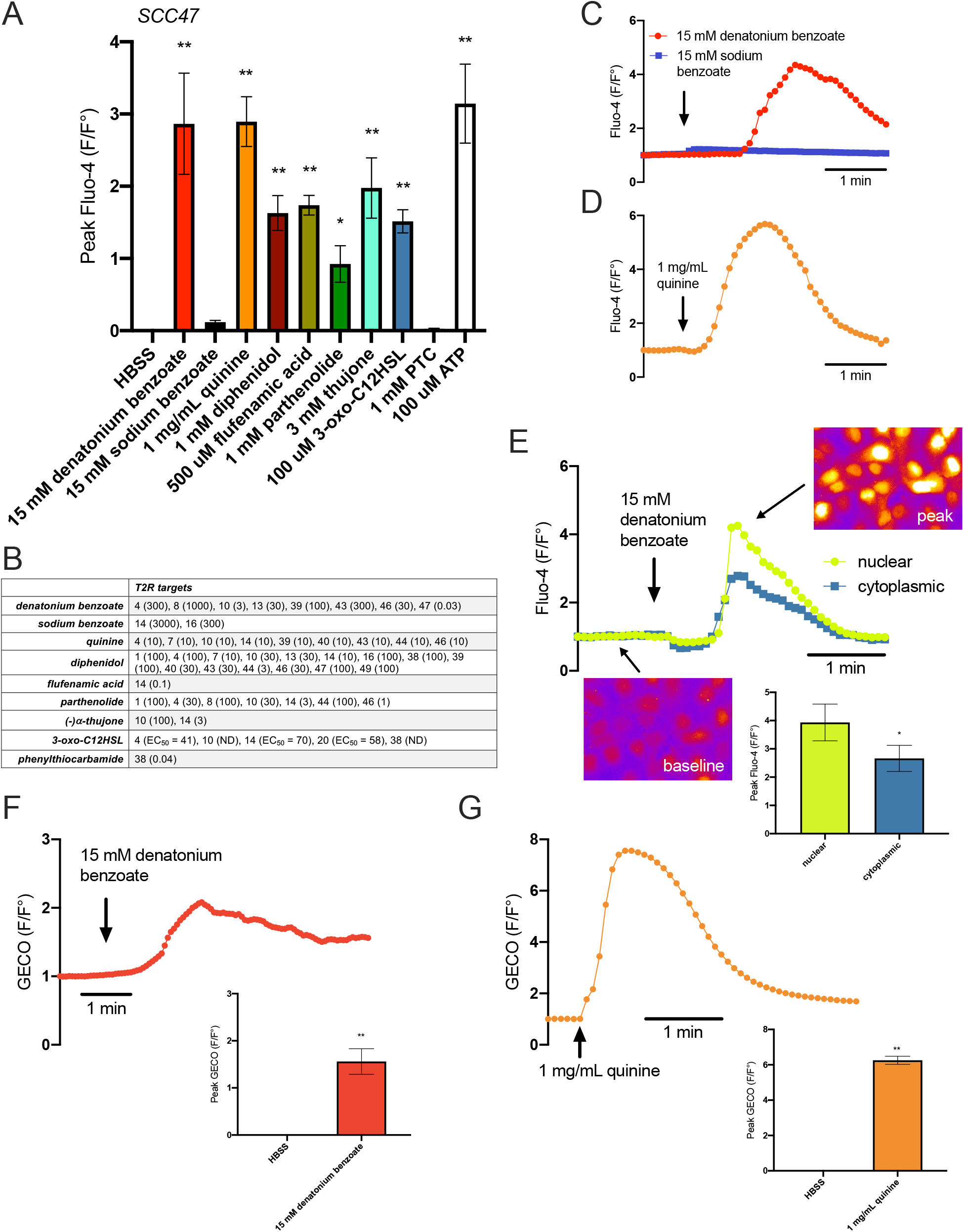
Bitter (T2R) agonists activate intracellular calcium (Ca^2+^_i_) responses in cultured head and neck squamous cell carcinoma (HNSCC). HNSCC cell line SCC47 loaded with Ca^2+^ binding dye, Fluo-4, was stimulated with T2R agonists and Ca^2+^ was measured over time. **A** Peak Fluo-4 F/F_o_ was quantified after stimulation with Hank’s Balanced Salt Solution (HBSS), denatonium benzoate, sodium benzoate, quinine, diphenidol, flufenamic acid, parthenolide, thujone, N-3-oxo-dodecanoyl-L-homoserine lactone (3-oxo-C12HSL), phenylthiocarbamide (PTC), and purinergic receptor agonist adenosine triphosphate (ATP) (mean ± SEM; 3-7 experiments using separate cultures). Significance by 1-way ANOVA with Bonferroni post-test comparing HBSS to each agonist. **B** Table of T2R targets for agonists used. Effective concentration (EC, in µM) or half maximal effective concentration (EC_50_, when indicated) shown in parentheses, taken from [37, 47]. Compounds with “ND” denote EC not determined. **C-D** Representative traces of Fluo-4 F/F_o_ over time after stimulation with denatonium benzoate (C) and sodium benzoate (separate traces superimposed) and quinine (D). **E** Fluo-4 fluorescence in response to denatonium benzoate-induced Ca^2+^ release appears to localize more strongly to the nucleus than the cytoplasm as represented by the traces and images; peak Fluo-4 F/F_o_ for nuclear Ca^2+^ release is significantly greater than cytoplasmic Ca^2+^ release (mean ± SEM; 3 separate cultures). Significance by unpaired t-test. **F-G** Nuclear-localized R-GECO-nls was used to measure nuclear Ca^2+^ after stimulation with (F) denatonium benzoate and (G) quinine. Both agonists stimulate nuclear Ca^2+^ as demonstrated by the traces over time and peak GECO F/F_o_ compared to HBSS (mean ± SEM; 3 separate cultures). Significance by unpaired t-test. \**p*<0.05; \*\**p*<0.01.

In heterologous expression studies, denatonium benzoate activates ∼8 T2Rs (**Fig. 3B**) while sodium benzoate activates only T2R14 and T2R16 very weakly (mM effective concentrations) [37] (**Fig. 3B**). Therefore, we used sodium benzoate as a pH and osmolarity control. Denatonium benzoate activated Ca^2+^_i_ responses while equimolar sodium benzoate did not (**Fig. 3A, C**), suggesting that the observed Ca^2+^_i_ is likely due to activation of specific T2Rs via the denatonium moiety.

Fitting with GPCR activation, denatonium benzoate and thujone Ca^2+^_i_ responses in SCC4 and SCC47 cells were inhibited by phospholipase C (PLC) inhibitor U73122 (**Supplementary Fig. S4A-C**). Ca^2+^_i_ responses to denatonium benzoate, thujone, and *P. aeruginosa* 3-oxo-C12HSL were also blocked by heterotrimeric G protein inhibitor YM254890 [38] (**Supplementary Fig. S4D-G**). Consistent with GPCR-induced PLC activation, Ca^2+^_i_ responses to denatonium benzoate and quinine were also blocked with inositol trisphosphate (IP_3_) receptor (IP_3_R) inhibitor xestospongin C (**Supplementary Fig. S4H-J**).

Fluo-4 responses to denatonium benzoate appeared primarily nuclear (shown for SCC47 in **Fig. 3E)**. This is similar to our studies of airway squamous cells showing T2Rs primarily regulate nuclear Ca^2+^ (Ca^2+^_nuc_) (preprint [36]). To determine if bitter agonists elevate Ca^2+^_nuc_, SCC47 cells were transfected with nuclear-localized genetically encoded calcium biosensor (R-GECO-nls [32]). Denatonium benzoate (15 mM) and quinine (1 mg/mL) triggered Ca^2+^_nuc_ elevation (**Fig. 3F, G**). Thus, T2Rs in HNSCC cells respond to a wide range of bitter agonists to trigger intracellular and intranuclear Ca^2+^ via GPCR and PLC.

### Nuclear Ca^2+^ leads to mitochondrial dysfunction and apoptosis

In some cells loaded with Fluo-8, we visualized an initial Ca^2+^_nuc_ response that was followed by a sustained peri-nuclear response that appeared mitochondrial (**Supplementary Fig. S5**). This led us to investigate how bitter agonists affect mitochondrial function in HNSCC cells. Ca^2+^_nuc_ may feed Ca^2+^ to the mitochondria and regulate metabolism or apoptosis [39]. We loaded SCC4 and 47 cells with ratiometric mitochondrial membrane potential (ΔΨ_m_) dye JC-1 and exposed them to denatonium benzoate or quinine over 6 hours. Both depolarized ΔΨ_m_, evidenced by a shift in the green-to-red fluorescence ratio (**Fig. 4A-D**). We also tested ΔΨ_m_ using tetramethylrhodamine ethyl ester (TMRE), which binds only to mitochondria with intact membrane potential. SCC4 cells were treated with denatonium benzoate and quinine for different lengths of time, stained with TMRE and Red Dot 2, and fluorescence intensities were compared (**Fig. 4E-H**). Both bitter agonists led to mitochondrial depolarization and subsequent plasma membrane permeability.

**Fig. 4.**
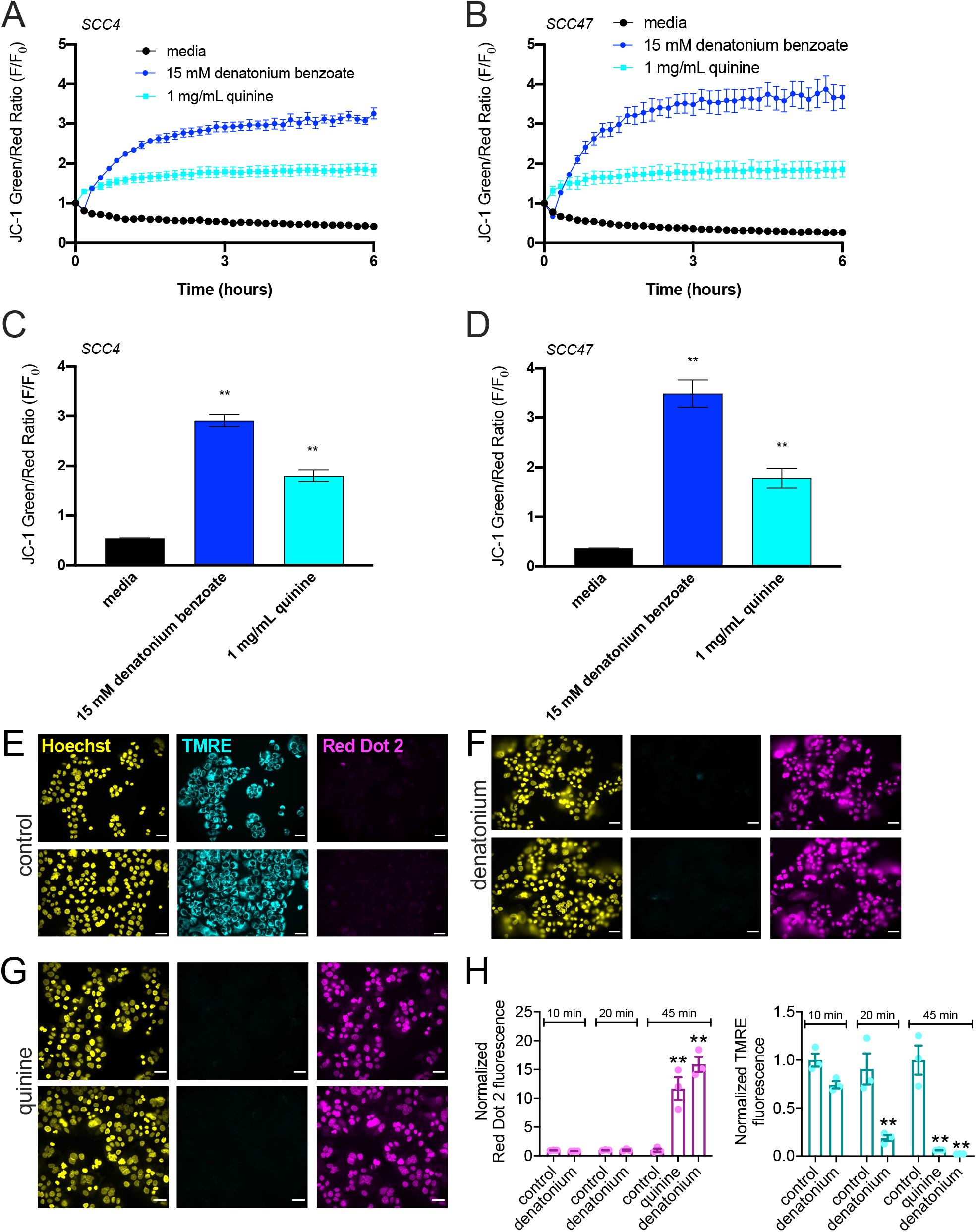
Bitter (T2R) agonists cause mitochondrial depolarization of head and neck squamous cell carcinoma (HNSCC). JC-1 dye, an indicator of mitochondrial membrane potential (ΔΨ_m_), shows higher green/red signal with depolarized mitochondria. **A-D** SCC cell lines loaded with JC-1 were stimulated with T2R agonists. Representative traces over 6 hours for HNSCC cell lines SCC4 (A) and SCC47 (B) after stimulation with media, denatonium benzoate, and quinine demonstrate mitochondrial depolarization with bitter agonists (mean ± SEM; 3 separate cultures for each cell line). JC-1 green/red ratio was quantified for SCC4 (C) and SCC47 (D) at 3 hours showing significantly higher ratios for denatonium benzoate and quinine compared to media (mean ± SEM; 3 separate cultures for each cell line). Significance by 1-way ANOVA with Bonferroni post-test comparing media to each agonist. **E-H** SCC4 was treated with media (E), denatonium benzoate (F), and quinine (G) for 45 minutes then stained with Hoechst (nuclei), TMRE (mitochondria with intact membrane potential), and Red Dot 2 (indicates permeable plasma membrane) dyes. Scale bars in *E-G* are 20 µm. **H** TMRE and Red Dot 2 fluorescence intensities were separately compared between media, denatonium benzoate, and quinine, demonstrating mitochondrial depolarization and plasma membrane permeability after bitter agonist stimulation (mean ± SEM; 3 separate cultures). Significance by 1-way ANOVA with Dunnett’s post-test comparing media to each agonist. \*\**p*<0.01.

Bitter agonists also decreased cellular metabolism, evidenced by a reduction in NAD(P)H production measured by XTT. In the presence of a cofactor, extracellular XTT is reduced from a non-light-absorbing to a colored absorbing form by cellular NAD(P)H via cross-plasma-membrane electron transfer. We saw blunted XTT changes 3 hours in the presence of 1-10 mM denatonium benzoate in SCC4 and 5-10 mM denatonium benzoate in SCC47 cells. Primary oral keratinocytes did not exhibit significant changes in XTT at 5-10 mM denatonium benzoate (**Fig. 5**).

**Fig. 5.**
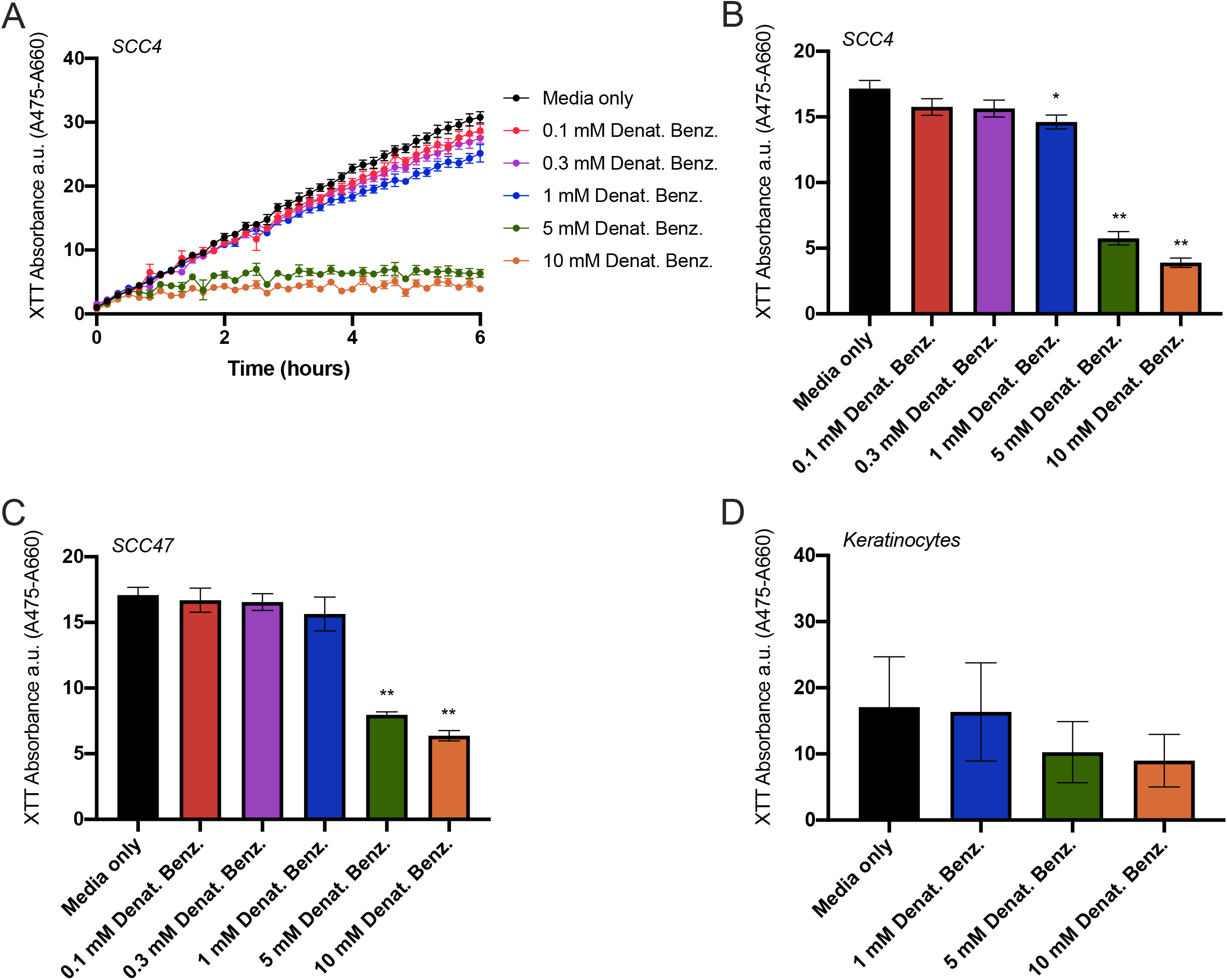
Bitter (T2R) agonists decrease cellular metabolism in head and neck squamous cell carcinoma (HNSCC). 2,3-bis-(2-methoxy-4-nitro-5-sulfophenyl)-2H-tetrazolium-5-carboxanilide (XTT) was used to measure cellular NAD(P)H production via a cross-plasma-membrane electron transfer. **A** Representative trace over 6 hours after stimulation of HNSCC cell line SCC4 with media ± denatonium benzoate at five different concentrations as indicated. **B-C** Bar graphs (mean ± SEM; n=6 independent cultures per cell line) of XTT absorbance quantified at 3 hours for **B** SCC4 and **C** SCC47. **D** XTT absorbance quantified at 3 hours for primary oral keratinocytes showing smaller (not statistically significant) reduction in NAD(P)H production with denatonium compared with HNSCC cells (n=6 independent cultures from different patients). Significance by 1-way ANOVA with Dunnett’s post-test comparing media to each concentration of denatonium benzoate. \**p*<0.05, \*\**p*<0.01.

To test if mitochondrial impairment led to apoptosis, we treated HNSCC cells with a caspase 3/7-sensitive dye (CellEvent caspase 3/7 reagent, Thermo). Exposure to denatonium benzoate caused a significant increase in caspase 3/7 activity over 6 hours in SCC4, SCC15, SCC47, and SCC152. Quinine led to a significant increase in SCC4, SCC15, and SCC47, but not SCC152 (**Fig. 6**). Denatonium benzoate but not sodium benzoate-induced caspase activation was confirmed using an optical assay (Flip-GFP; [40]) in SCC4, SCC47, SCC90, and SCC152 cells (**Supplementary Fig. S6-7**).

**Fig. 6.**
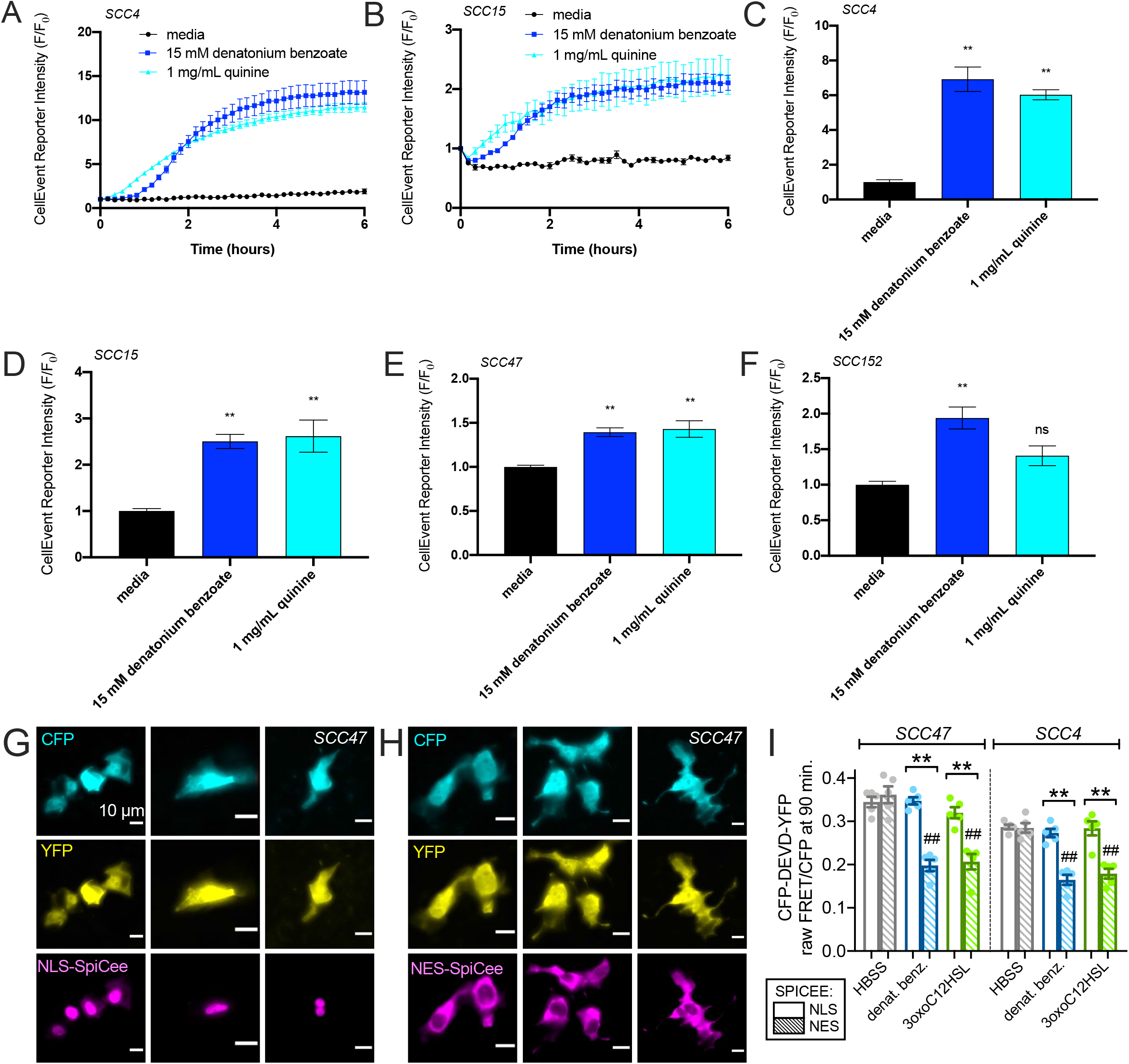
Bitter (T2R) agonists cause activation of apoptosis in head and neck squamous cell carcinoma (HNSCC). **A-B** Cleavage of caspase 3/7 CellEvent reagent signals apoptosis. Representative traces over 6 hours after stimulation of HNSCC cell lines SCC4 (A) and SCC15 (B) with media, denatonium benzoate, and quinine demonstrate increased caspase 3/7 cleavage (mean ± SEM; 3-5 separate cultures for each cell line). **C-F** CellEvent reporter intensity at 6 hours after stimulation of SCC4 (C), SCC15 (D), SCC47 (E), and SCC152 (F) with bitter agonists, normalized to media (mean ± SEM; 3-5 separate cultures for each cell line). Significance by 1-way ANOVA with Bonferroni post-test comparing media to each agonist; \*\**p*<0.01, ns=non-significant. **G-H** Wide-field 40x fluorescence images of mCherry-labeled NLS-SpiCee (G) or NES-SpiCee (H) co-transfected with caspase FRET biosensor in SCC47 cells. **I** FRET ratios were measured in cells after 90 min incubation with HBSS (gray bars), 10 mM denatonium benzoate (blue bars), or 100 µM 3-oxo-C12HSL (green bars). Open bars are NLS-SpiCee and crossed bars are NES-SpiCee. A downward deflection indicates loss of FRET signal signifying caspase cleavage. Each data point is an independent experiment. Each experiment images 5-9 cells from a single field of view. Significance by one-way ANOVA with Bonferonni posttest; \*\**p*<0.01 between bracketed bars; ^##^p<0.01 compared with HBSS.

We further confirmed caspase activation in SCC4 and SCC47 cells using a Förester resonance energy transfer (FRET)-based biosensor created by linking enhanced cyan fluorescent protein (eCFP) and yellow fluorescent protein variant mVenus with a linker containing a DEVD caspase 3 cleavage site [41]. Caspase cleavage of the protein allows the eCFP and mVenus to diffuse farther apart, reducing FRET (**Supplementary Fig. S8A**). We saw FRET decreases signaling caspase activation in response to denatonium benzoate, quinine, thujone, 3-oxo-C12HSL, and *Pseudomonas* quinolone signal (PQS) but not sodium benzoate (**Supplementary Fig. S8B-F**).

Loading SCC4 or SCC47 cells with calcium chelator BAPTA blocked caspase activation in response to denatonium benzoate (**Supplementary Fig. S6**). To determine if apoptosis was linked to Ca^2+^_nuc_, we utilized a recombinant protein calcium buffer, known as SpiCee [42], fused to either nuclear export or nuclear localization sequences (NES-SpiCee or NLS-SpiCee, respectively). NLS-SpiCee or NES-SpiCee was co-transfected with the eCFP-DEVD-Venus biosensor (**Fig. 6G-H**). FRET decreases signaling apoptosis occured with NES-SpiCee and bitter agonists denatonium benzoate or bacterial 3-oxo-C12HSL (**Fig. 6I**). However, no apoptosis was observed with NLS-SpiCee. Chelation of Ca^2+^_nuc_ blocked bitter agonist-induced apoptosis (**Fig. 6I**). Bitter agonist-induced Ca^2+^_nuc_ responses reduce cellular metabolism and proliferation in HNSCC cells.

### Increased *TAS2R* expression may be associated with HNSCC survival

Data above suggest T2Rs activate apoptosis and limit proliferation of HNSCCs *in vitro*, which prompted us to explore possible *in vivo* effects of T2Rs. Specifically, we investigated the impact of increased *TAS2R* expression on survival in HNSCC using TCGA [34, 35]. Analysis included 504 cases of HNSCC diagnosed between 1992 and 2013 with mRNA expression data available. *TAS2R* expression z-scores relative to diploid samples are demonstrated in the heatmap in **Fig. 7A**. Kaplan-Meier survival analysis of cases with high vs low *TAS2R* expression demonstrated improved overall survival for cases with increased *TAS2R* expression (p = 0.0208 by logrank test, **Fig. 7B**). Median survival was 65.77 months for high *TAS2R* expression versus 39.49 months for low mRNA expression. We also separately evaluated *TAS2R4* expression, as this receptor was expressed in the patient samples and cell lines and is activated by denatonium benzoate and quinine. Kaplan-Meier survival analysis comparing high *TAS2R4* expression and low expression demonstrated improved overall survival for cases with increased *TAS2R4* (p = 0.0269 by logrank test, **Fig. 7C**).

**Fig. 7.**
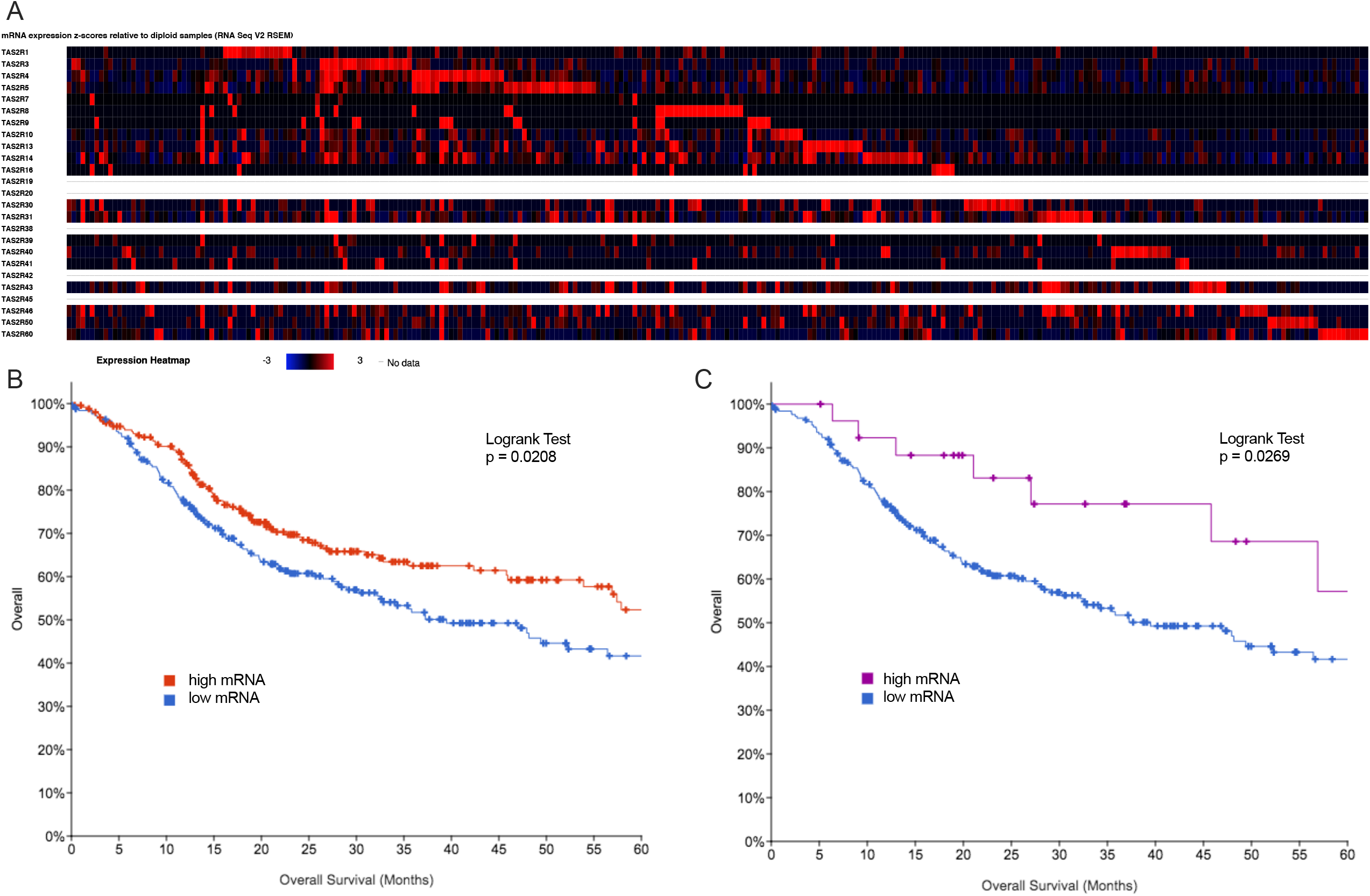
Bitter taste receptor (*TAS2R*) expression alterations in head and neck squamous cell carcinoma (HNSCC) are associated with survival. *TAS2R* expression alterations were analyzed for 504 cases of HNSCC using The Cancer Genome Atlas (TCGA) [34, 35]. **A** Heatmap of *TAS2R* mRNA expression z-scores relative to diploid samples. Increased *TAS2R* expression (red) is more common than decreased expression (blue). **B** 5-year survival analysis of HNSCC cases demonstrate that increased *TAS2R* expression is associated with improved overall survival (p = 0.0208 by logrank test). **C** Increased *TAS2R*4 expression, which is activated by the bitter agonists denatonium benzoate, quinine, diphenidol, parthenolide, and 3-oxo-C12HSL, is also associated with improved overall survival (p = 0.0269 by logrank test).

Evaluation of genetic changes in addition to expression changes showed that 283 out of 504 cases (56.15%) had some *TAS2R* alteration (**Supplementary Fig. S9A**). Multiple alterations were present in 11.51% of cases (**Supplementary Fig. S9B**). TCGA analyses suggest *TAS2R* genetic and expression alterations are prevalent in HNSCC. Increased *TAS2R* expression may be beneficial by limiting cancer cell proliferation via mechanisms outlined above.

## Discussion

While T2Rs have been studied in cancer [14-23], this is the first description of T2R expression and signaling in HNSCC. Bitter agonists activate T2R-mediated Ca^2+^_nuc_, which triggers mitochondrial depolarization, caspase activation, and apoptosis. The novel demonstration of a requirement for Ca^2+^_nuc_ in bitter agonist-induced apoptosis may be important to other cancers. Using TCGA, we report that increased *TAS2R* expression is correlated with HNSCC survival, suggesting potential clinical utility of T2R agonists and supporting future prospective studies in larger patient populations to test if/how T2Rs are diagnostic biomarkers in HNSCC.

The *TAS2Rs* with highest expression in HNSCC (*TAS2R4, 14, 19, 20, 30, 43, 45*) overlap with *TAS2Rs* expressed at increased levels in breast cancer [20, 21]. However, the concept of T2R expression on the nuclear membrane is highly novel. We (preprint [36]) and others [43] have found nuclear localization of T2Rs in de-ciliated or inflamed airway tissues. We also saw this in lung cancer cells (preprint [36]). Squamous de-differentiation likely alters localization of airway T2Rs from cilia to the nucleus, where T2Rs activate Ca^2+^_nuc_ and apoptosis rather than ciliary beating activated in differentiated healthy tissues [10, 33]. HNSCC cells also have at least some nuclear T2Rs. Prior studies have validated importance of nuclear envelope GPCRs in neurons and other cells [44-46], but the link between intracellular T2Rs, Ca^2+^_nuc_, and apoptosis in cancer cells is novel.

Interestingly, orphan T2R42 [37, 47] localized to the nucleus in two HNSCC tongue cell lines. Several T2Rs that are activated by denatonium benzoate or quinine (including T2Rs 4, 8, 10, 14, 30/47, and 46) did not localize to the nucleus, but nonetheless appeared intracellular. T2R13, activated by denatonium benzoate [37, 47], was nuclear in SCC47 but not in SCC4. Thus, differences in T2R localization between cancers of the same anatomic site may exist.

Multi-T2R targeting compounds like denatonium benzoate or quinine might be useful to activate apoptosis in HNSCCs from a broad range of patients. Many known T2R agonists (e.g., quinine) are cell permeant [48, 49]. Intracellular T2Rs could be targeted using high-dose topical treatments in accessible anatomic locations like the oral cavity/oropharynx. Many pharmaceuticals with known safety profiles are bitter [50], and could potentially kill or slow the growth of HNSCC, particularly if specific T2R agonists can be used in combination with genetic profiling to determine expression of specific T2Rs in tumor vs normal tissue.

Ca^2+^_nuc_ and cytosolic Ca^2+^ are distinct, but both are activated by ryanodine receptors and IP_3_Rs [51, 52]. Ca^2+^_nuc_ responses to bitter agonists in HNSCC originates partly from the ER, requiring PLC and IP_3_Rs. Nuclear Ca^2+^ regulates transcription factors but was also linked to apoptosis [53-55]. Bitter agonists activate apoptosis and/or mitochondrial depolarization in other cell types [56-59], including metastatic breast cancer [60], prostate cancer [61], and acute myeloid leukemia cells [59]. In pancreatic cancer, T2R38 can be activated by 3-oxo-C12HSL [15], suggesting a possible link with the microbiome. In contrast, denatonium benzoate had pro-tumor actions in murine submandibular gland cancer cells [23]. We show here that bitter agonists, including bacterial 3-oxo-C12HSL and PQS, are anti-proliferative and pro-apoptotic in HNSCC, with a novel mechanistic link between Ca^2+^_nuc_ and apoptosis.

TCGA analysis suggests *TAS2R* expression alterations impact HNSCC or serve as markers of underlying genetic changes that contribute to tumor biology and treatment efficacy. *TAS2R* expression alterations are associated with better survival. We hypothesize that, as T2Rs regulate apoptosis, tumors that overexpress T2Rs may have an improved prognosis due to a more robust internal regulation preventing unchecked proliferation or responding to bitter agonists in the microenvironment, either from the tumor microbiome or intracellular metabolites. Further work is necessary to appreciate the full impact of T2Rs in HNSCC and potential roles in risk stratification and treatment. Furthermore, common genetic polymorphisms in *TAS2Rs* that affect bitter taste perception and dietary preferences [62, 63] may contribute to differences in tumor behavior. Importantly, bacterial ligands like PQS or 3-oxo-C12HSL are hydrophobic and cell-permeant [64, 65]. We propose that cancer cell T2Rs link the host microbiome with HNSCC progression. HNSCCs are known to be associated with unique microbiomes [66-70], impacting the immune system and contributing to oncogenesis and oncologic outcomes [66, 71, 72].

Identification of prognostic markers to guide treatment selection is an active area of HNSCC research to ultimately improve survival while mitigating treatment-related side effects. Several clinical trials have sought to limit treatment-related morbidity by de-intensifying therapy in HPV-associated oropharyngeal tumors, which are known to have improved prognosis compared to HPV-negative tumors [73, 74]. If increased *TAS2R* expression portends favorable prognoses, it would be reasonable to perform *TAS2R* expression analysis of HNSCC tumors as part of the treatment selection algorithm, similar to standard practice HPV testing for oropharyngeal SCCs.

### Materials availability

All materials will be provided under Materials Transfer Agreement upon request to RJL.

## Acknowledgements

We thank Maureen Victoria (University of Pennsylvania) for technical assistance and Noam Cohen (University of Pennsylvania) for advice and discussion. We thank Dr. Hans Joenje (VU Medical Center, Netherlands) for providing VU147T cells.

## Author contributions

Conceptualization and Visualization: R.M.C., E.A.W., R.J.L.; Investigation and Formal Analysis: R.M.C., D.B.M., I.G., and R.J.L.; Writing – Original Draft: R.M.C.; Writing – Review and Editing: R.M.C., E.A.W., and R.J.L.; Data Curation and Resources: K.R, J.G.N., and D.B.; Funding Acquisition: R.M.C. and R.J.L.; Supervision: R.J.L.

## Ethical approval

Tissue collection was approved by University of Pennsylvania Institutional Review Board (protocol #417200)

## REFERENCES

1. Johnson DE, Burtness B, Leemans CR, Lui VWY, Bauman JE, Grandis JR. Head and neck squamous cell carcinoma. Nature reviews Disease primers 2020, 6(1): 92.

2. Jou A, Hess J. Epidemiology and Molecular Biology of Head and Neck Cancer. Oncol Res Treat 2017, 40(6): 328–332.

3. Kamangar F, Dores GM, Anderson WF. Patterns of cancer incidence, mortality, and prevalence across five continents: defining priorities to reduce cancer disparities in different geographic regions of the world. J Clin Oncol 2006, 24(14): 2137–2150.

4. Network NCC. Head and Neck Cancers (Version 2.2019). 2019 [cited August 16, 2019]Available from: https://www.nccn.org/professionals/physician_gls/pdf/head-and-neck.pdf

5. Cramer JD, Burtness B, Ferris RL. Immunotherapy for head and neck cancer: Recent advances and future directions. Oral Oncol 2019, 99: 104460.

6. Lazarus CL. Effects of chemoradiotherapy on voice and swallowing. Curr Opin Otolaryngol Head Neck Surg 2009, 17(3): 172–178.

7. Weinberger PM, Yu Z, Haffty BG, Kowalski D, Harigopal M, Brandsma J, et al. Molecular classification identifies a subset of human papillomavirus--associated oropharyngeal cancers with favorable prognosis. J Clin Oncol 2006, 24(5): 736–747.

8. Ang KK, Harris J, Wheeler R, Weber R, Rosenthal DI, Nguyen-Tan PF, et al. Human papillomavirus and survival of patients with oropharyngeal cancer. N Engl J Med 2010, 363(1): 24–35.

9. Janssen S, Laermans J, Verhulst PJ, Thijs T, Tack J, Depoortere I. Bitter taste receptors and alpha-gustducin regulate the secretion of ghrelin with functional effects on food intake and gastric emptying. Proceedings of the National Academy of Sciences of the United States of America 2011, 108(5): 2094–2099.

10. Hariri BM, McMahon DB, Chen B, Freund JR, Mansfield CJ, Doghramji LJ, et al. Flavones modulate respiratory epithelial innate immunity: anti-inflammatory effects and activation of the T2R14 receptor. J Biol Chem 2017, 292(20): 8484–8497.

11. Carey RM, Lee RJ. Taste Receptors in Upper Airway Innate Immunity. Nutrients 2019, 11(9).

12. Bachmanov AA, Bosak NP, Lin C, Matsumoto I, Ohmoto M, Reed DR, et al. Genetics of taste receptors. Current pharmaceutical design 2014, 20(16): 2669–2683.

13. Gopallawa I, Freund JR, Lee RJ. Bitter taste receptors stimulate phagocytosis in human macrophages through calcium, nitric oxide, and cyclic-GMP signaling. Cellular and molecular life sciences : CMLS 2020.

14. Stern L, Giese N, Hackert T, Strobel O, Schirmacher P, Felix K, et al. Overcoming chemoresistance in pancreatic cancer cells: role of the bitter taste receptor T2R10. Journal of Cancer 2018, 9(4): 711–725.

15. Gaida MM, Mayer C, Dapunt U, Stegmaier S, Schirmacher P, Wabnitz GH, et al. Expression of the bitter receptor T2R38 in pancreatic cancer: localization in lipid droplets and activation by a bacteria-derived quorum-sensing molecule. Oncotarget 2016, 7(11): 12623–12632.

16. Choi JH, Lee J, Choi IJ, Kim YW, Ryu KW, Kim J. Genetic Variation in the TAS2R38 Bitter Taste Receptor and Gastric Cancer Risk in Koreans. Scientific reports 2016, 6: 26904.

17. Yamaki M, Saito H, Isono K, Goto T, Shirakawa H, Shoji N, et al. Genotyping Analysis of Bitter-Taste Receptor Genes TAS2R38 and TAS2R46 in Japanese Patients with Gastrointestinal Cancers. Journal of nutritional science and vitaminology 2017, 63(2): 148–154.

18. Carrai M, Steinke V, Vodicka P, Pardini B, Rahner N, Holinski-Feder E, et al. Association between TAS2R38 gene polymorphisms and colorectal cancer risk: a case-control study in two independent populations of Caucasian origin. PloS one 2011, 6(6): e20464.

19. Barontini J, Antinucci M, Tofanelli S, Cammalleri M, Dal Monte M, Gemignani F, et al. Association between polymorphisms of TAS2R16 and susceptibility to colorectal cancer. BMC gastroenterology 2017, 17(1): 104.

20. Jaggupilli A, Singh N, Upadhyaya J, Sikarwar AS, Arakawa M, Dakshinamurti S, et al. Analysis of the expression of human bitter taste receptors in extraoral tissues. Mol Cell Biochem 2017, 426(1-2): 137–147.

21. Singh N, Chakraborty R, Bhullar RP, Chelikani P. Differential expression of bitter taste receptors in non-cancerous breast epithelial and breast cancer cells. Biochemical and biophysical research communications 2014, 446(2): 499–503.

22. Choi JH, Lee J, Yang S, Lee EK, Hwangbo Y, Kim J. Genetic variations in TAS2R3 and TAS2R4 bitterness receptors modify papillary carcinoma risk and thyroid function in Korean females. Scientific reports 2018, 8(1): 15004.

23. Dmytrenko G, Castro ME, Sales ME. Denatonium and Naringenin Promote SCA-9 Tumor Growth and Angiogenesis: Participation of Arginase. Nutrition and cancer 2017, 69(5): 780–790.

24. Lambert JD, VanDusen SR, Cockroft JE, Smith EC, Greenwood DC, Cade JE. Bitter taste sensitivity, food intake, and risk of malignant cancer in the UK Women’s Cohort Study. European journal of nutrition 2019, 58(5): 2111–2121.

25. Sriram K, Moyung K, Corriden R, Carter H, Insel PA. GPCRs show widespread differential mRNA expression and frequent mutation and copy number variation in solid tumors. PLoS Biol 2019, 17(11): e3000434.

26. McMahon DB, Carey RM, Kohanski MA, Tong CCL, Papagiannopoulos P, Adappa ND, et al. Neuropeptide regulation of secretion and inflammation in human airway gland serous cells. Eur Respir J 2020, 55(4).

27. McMahon DB, Workman AD, Kohanski MA, Carey RM, Freund JR, Hariri BM, et al. Protease-activated receptor 2 activates airway apical membrane chloride permeability and increases ciliary beating. FASEB J 2018, 32(1): 155–167.

28. Carey RM, Freund JR, Hariri BM, Adappa ND, Palmer JN, Lee RJ. Polarization of protease-activated receptor 2 (PAR-2) signaling is altered during airway epithelial remodeling and deciliation. J Biol Chem 2020, 295(19): 6721–6740.

29. McMahon DB, Carey RM, Kohanski MA, Adappa ND, Palmer JN, Lee RJ. PAR-2-activated secretion by airway gland serous cells: role for CFTR and inhibition by Pseudomonas aeruginosa. Am J Physiol Lung Cell Mol Physiol 2021, 320(5): L845–L879.

30. Basu D, Nguyen TT, Montone KT, Zhang G, Wang LP, Diehl JA, et al. Evidence for mesenchymal-like sub-populations within squamous cell carcinomas possessing chemoresistance and phenotypic plasticity. Oncogene 2010, 29(29): 4170–4182.

31. Begum S, Gillison ML, Ansari-Lari MA, Shah K, Westra WH. Detection of human papillomavirus in cervical lymph nodes: a highly effective strategy for localizing site of tumor origin. Clin Cancer Res 2003, 9(17): 6469–6475.

32. Zhao Y, Araki S, Wu J, Teramoto T, Chang YF, Nakano M, et al. An expanded palette of genetically encoded Ca(2)(+) indicators. Science 2011, 333(6051): 1888–1891.

33. Freund JR, Mansfield CJ, Doghramji LJ, Adappa ND, Palmer JN, Kennedy DW, et al. Activation of airway epithelial bitter taste receptors by Pseudomonas aeruginosa quinolones modulates calcium, cyclic-AMP, and nitric oxide signaling. J Biol Chem 2018, 293(25): 9824–9840.

34. Cerami E, Gao J, Dogrusoz U, Gross BE, Sumer SO, Aksoy BA, et al. The cBio cancer genomics portal: an open platform for exploring multidimensional cancer genomics data. Cancer discovery 2012, 2(5): 401–404.

35. Gao J, Aksoy BA, Dogrusoz U, Dresdner G, Gross B, Sumer SO, et al. Integrative analysis of complex cancer genomics and clinical profiles using the cBioPortal. Sci Signal 2013, 6(269): pl1.

36. McMahon DB, Kuek LE, Johnson ME, Johnson PO, Horn RLJ, Carey RM, et al. The bitter end: T2R bitter receptor agonists elevate nuclear calcium and induce apoptosis in non-ciliated airway epithelial cells. bioRxiv 2021: 2021.2005.2016.444376.

37. Meyerhof W, Batram C, Kuhn C, Brockhoff A, Chudoba E, Bufe B, et al. The molecular receptive ranges of human TAS2R bitter taste receptors. Chem Senses 2010, 35(2): 157–170.

38. Tietze D, Kaufmann D, Tietze AA, Voll A, Reher R, Konig G, et al. Structural and Dynamical Basis of G Protein Inhibition by YM-254890 and FR900359: An Inhibitor in Action. J Chem Inf Model 2019, 59(10): 4361–4373.

39. Nicotera P, Zhivotovsky B, Orrenius S. Nuclear calcium transport and the role of calcium in apoptosis. Cell Calcium 1994, 16(4): 279–288.

40. Zhang Q, Schepis A, Huang H, Yang J, Ma W, Torra J, et al. Designing a Green Fluorogenic Protease Reporter by Flipping a Beta Strand of GFP for Imaging Apoptosis in Animals. J Am Chem Soc 2019, 141(11): 4526–4530.

41. Albeck JG, Burke JM, Spencer SL, Lauffenburger DA, Sorger PK. Modeling a snap-action, variable-delay switch controlling extrinsic cell death. PLoS Biol 2008, 6(12): 2831–2852.

42. Ros O, Baudet S, Zagar Y, Loulier K, Roche F, Couvet S, et al. SpiCee: A Genetic Tool for Subcellular and Cell-Specific Calcium Manipulation. Cell Rep 2020, 32(3): 107934.

43. Zborowska-Piskadlo K, Stachowiak M, Rusetska N, Sarnowska E, Siedlecki J, Dzaman K. The expression of bitter taste receptor TAS2R38 in patients with chronic rhinosinusitis. Arch Immunol Ther Exp (Warsz) 2020, 68(5): 26.

44. Jong YI, Harmon SK, O’Malley KL. GPCR signalling from within the cell. Br J Pharmacol 2018, 175(21): 4026–4035.

45. Crilly SE, Puthenveedu MA. Compartmentalized GPCR Signaling from Intracellular Membranes. J Membr Biol 2020.

46. Ribeiro-Oliveira R, Vojtek M, Goncalves-Monteiro S, Vieira-Rocha MS, Sousa JB, Goncalves J, et al. Nuclear G-protein-coupled receptors as putative novel pharmacological targets. Drug Discov Today 2019, 24(11): 2192–2201.

47. Wiener A, Shudler M, Levit A, Niv MY. BitterDB: a database of bitter compounds. Nucleic acids research 2012, 40(Database issue): D413–419.

48. Peri I, Mamrud-Brains H, Rodin S, Krizhanovsky V, Shai Y, Nir S, et al. Rapid entry of bitter and sweet tastants into liposomes and taste cells: implications for signal transduction. Am J Physiol Cell Physiol 2000, 278(1): C17–25.

49. Zubare-Samuelov M, Shaul ME, Peri I, Aliluiko A, Tirosh O, Naim M. Inhibition of signal termination-related kinases by membrane-permeant bitter and sweet tastants: potential role in taste signal termination. Am J Physiol Cell Physiol 2005, 289(2): C483–492.

50. Levit A, Nowak S, Peters M, Wiener A, Meyerhof W, Behrens M, et al. The bitter pill: clinical drugs that activate the human bitter taste receptor TAS2R14. FASEB J 2014, 28(3): 1181–1197.

51. Gomes DA, Leite MF, Bennett AM, Nathanson MH. Calcium signaling in the nucleus. Can J Physiol Pharmacol 2006, 84(3-4): 325–332.

52. Rodrigues MA, Gomes DA, Nathanson MH, Leite MF. Nuclear calcium signaling: a cell within a cell. Braz J Med Biol Res 2009, 42(1): 17–20.

53. Gerasimenko OV, Gerasimenko JV, Tepikin AV, Petersen OH. Calcium transport pathways in the nucleus. Pflugers Arch 1996, 432(1): 1–6.

54. Petersen OH, Gerasimenko OV, Gerasimenko JV, Mogami H, Tepikin AV. The calcium store in the nuclear envelope. Cell Calcium 1998, 23(2-3): 87–90.

55. Martelli AM, Mazzotti G, Capitani S. Nuclear protein kinase C isoforms and apoptosis. Eur J Histochem 2004, 48(1): 89–94.

56. Wen X, Zhou J, Zhang D, Li J, Wang Q, Feng N, et al. Denatonium inhibits growth and induces apoptosis of airway epithelial cells through mitochondrial signaling pathways. Respir Res 2015, 16: 13.

57. Sharma P, Panebra A, Pera T, Tiegs BC, Hershfeld A, Kenyon LC, et al. Antimitogenic effect of bitter taste receptor agonists on airway smooth muscle cells. Am J Physiol Lung Cell Mol Physiol 2016, 310(4): L365–376.

58. Pan S, Sharma P, Shah SD, Deshpande DA. Bitter taste receptor agonists alter mitochondrial function and induce autophagy in airway smooth muscle cells. Am J Physiol Lung Cell Mol Physiol 2017, 313(1): L154–l165.

59. Salvestrini V, Ciciarello M, Pensato V, Simonetti G, Laginestra MA, Bruno S, et al. Denatonium as a Bitter Taste Receptor Agonist Modifies Transcriptomic Profile and Functions of Acute Myeloid Leukemia Cells. Frontiers in oncology 2020, 10: 1225.

60. Singh N, Shaik FA, Myal Y, Chelikani P. Chemosensory bitter taste receptors T2R4 and T2R14 activation attenuates proliferation and migration of breast cancer cells. Mol Cell Biochem 2020.

61. Martin LTP, Nachtigal MW, Selman T, Nguyen E, Salsman J, Dellaire G, et al. Bitter taste receptors are expressed in human epithelial ovarian and prostate cancers cells and noscapine stimulation impacts cell survival. Mol Cell Biochem 2019, 454(1–2): 203–214.

62. Kim UK, Jorgenson E, Coon H, Leppert M, Risch N, Drayna D. Positional cloning of the human quantitative trait locus underlying taste sensitivity to phenylthiocarbamide. Science 2003, 299(5610): 1221–1225.

63. Risso D, Tofanelli S, Morini G, Luiselli D, Drayna D. Genetic variation in taste receptor pseudogenes provides evidence for a dynamic role in human evolution. BMC Evol Biol 2014, 14: 198.

64. Schwarzer C, Fu Z, Patanwala M, Hum L, Lopez-Guzman M, Illek B, et al. Pseudomonas aeruginosa biofilm-associated homoserine lactone C12 rapidly activates apoptosis in airway epithelia. Cell Microbiol 2012, 14(5): 698–709.

65. Schwarzer C, Fu Z, Shuai S, Babbar S, Zhao G, Li C, et al. Pseudomonas aeruginosa homoserine lactone triggers apoptosis and Bak/Bax-independent release of mitochondrial cytochrome C in fibroblasts. Cell Microbiol 2014, 16(7): 1094–1104.

66. Banerjee S, Tian T, Wei Z, Peck KN, Shih N, Chalian AA, et al. Microbial Signatures Associated with Oropharyngeal and Oral Squamous Cell Carcinomas. Scientific reports 2017, 7(1): 4036.

67. Pushalkar S, Ji X, Li Y, Estilo C, Yegnanarayana R, Singh B, et al. Comparison of oral microbiota in tumor and non-tumor tissues of patients with oral squamous cell carcinoma. BMC Microbiol 2012, 12: 144.

68. Schmidt BL, Kuczynski J, Bhattacharya A, Huey B, Corby PM, Queiroz EL, et al. Changes in abundance of oral microbiota associated with oral cancer. PloS one 2014, 9(6): e98741.

69. Carey RM, Rajasekaran K, Seckar T, Lin X, Wei Z, Tong CCL, et al. The virome of HPV-positive tonsil squamous cell carcinoma and neck metastasis. Oncotarget 2020, 11(3): 282–293.

70. Rajasekaran K, Carey RM, Lin X, Seckar TD, Wei Z, Chorath K, et al. The microbiome of HPV-positive tonsil squamous cell carcinoma and neck metastasis. Oral Oncol 2021, 117: 105305.

71. Gopalakrishnan V, Helmink BA, Spencer CN, Reuben A, Wargo JA. The Influence of the Gut Microbiome on Cancer, Immunity, and Cancer Immunotherapy. Cancer Cell 2018, 33(4): 570–580.

72. Zitvogel L, Ma Y, Raoult D, Kroemer G, Gajewski TF. The microbiome in cancer immunotherapy: Diagnostic tools and therapeutic strategies. Science 2018, 359(6382): 1366–1370.

73. An Y, Holsinger FC, Husain ZA. De-intensification of adjuvant therapy in human papillomavirus-associated oropharyngeal cancer. Cancers Head Neck 2016, 1: 18.

74. Swisher-McClure S, Lukens JN, Aggarwal C, Ahn P, Basu D, Bauml JM, et al. A Phase 2 Trial of Alternative Volumes of Oropharyngeal Irradiation for De-intensification (AVOID): Omission of the Resected Primary Tumor Bed After Transoral Robotic Surgery for Human Papilloma Virus-Related Squamous Cell Carcinoma of the Oropharynx. Int J Radiat Oncol Biol Phys 2020, 106(4): 725–732.

