## Supplementary Material for "T2R bitter taste receptors regulate apoptosis and may be associated with survival in head and neck squamous cell carcinoma"

**Supplementary Table S1** Key biological and chemical reagents used in this study.

| REAGENT or RESOURCE | SOURCE | IDENTIFIER |
|---|---|---|
| Antibodies | | |
| anti-T2R8 | Abcam | ab75109 |
| anti-T2R13 | Abcam | ab172986 |
| anti-T2R4 | ThermoFisher Scientific | PA-67752 |
| anti-T2R10 | ThermoFisher Scientific | OSR00158W |
| anti-T2R14 | ThermoFisher Scientific | PA5-39710 |
| anti-T2R42 | ThermoFisher Scientific | NBP1-83154 |
| anti-T2R30/47 | ThermoFisher Scientific | PA5-67773 |
| anti-T2R46 | ThermoFisher Scientific | PA5-67772 |
| anti-Lamin B2 clone E-3 | ThermoFisher Scientific | 332100 |
| Alexa Fluor 488-conjugated donkey anti-mouse | ThermoFisher Scientific | A21202 |
| Alexa Fluor 546-conjugated donkey anti-rabbit | ThermoFisher Scientific | A10040 |
| anti-α-tubulin | Developmental Studies Hybridoma Bank | 12G10 |
| Chemical reagents | | |
| U73122 | Cayman Chemical | 70740 |
| U73343 | Cayman Chemical | 17339 |
| YM254890 | Cayman Chemical | 29735 |
| Xestospongin C | Cayman Chemical | 64950 |
| Parthenolide | Cayman Chemical | 70080 |
| Diphenhydramine | Cayman Chemical | 11158 |
| TMRE (Tetramethylrhodamine ethyl ester) | Cayman Chemical | 601283 |
| RedDot 2 Viability Dye | Cayman Chemical | 601282 |
| Cell-Based Hoechst Dye | Cayman Chemical | 600332 |
| ATP | Millipore Sigma | A9187 |
| (-)-a-Thujone | Millipore Sigma | 89231 |
| Flufenamic acid (FFA) | Millipore Sigma | F9005 |
| Denatonium benzoate | Millipore Sigma | D5765 |
| Phenylthiocarbamide (PTC) | Millipore Sigma | P7629 |
| Sodium Benzoate | Millipore Sigma | B3420 |
| Diphenidol | Millipore Sigma | SML2169 |
| Quinine | Millipore Sigma | Q0132 |
| N-3-oxo-dodecanoyl-L-homoserine lactone (3-oxo-C12HSL) | Millipore Sigma | O9139 |
| 2-Heptyl-3-hydroxy-4(1H)-quinolone (*Pseudomonas* quinolone signal, PQS) | Millipore Sigma | 94398 |
| Saponin | Millipore Sigma | S7900 |
| Bovine Serum Albumin | Millipore Sigma | A2153 |
| TRIzol | ThermoFisher Scientific | 15596026 |

| | | |
|---|---|---|
| JC-1 | ThermoFisher Scientific | T3168 |
| CellEvent™ Caspase-3/7 Green Detection Reagent | ThermoFisher Scientific | C10423 |
| XTT (sodium 3´-[1- (phenylaminocarbonyl)- 3,4-tetrazolium]-bis (4-methoxy6-nitro) benzene sulfonic acid hydrate | ThermoFisher Scientific | X6493 |
| lipofectamine 3000 | ThermoFisher Scientific | L3000075 |
| Fluo-4-AM | ThermoFisher Scientific | F14201 |
| Fluo-8-AM | Abcam | ab142773 |
| Normal Donkey Serum | Abcam | ab7475 |
| Critical Commercial Assays | | |
| High-Capacity cDNA Reverse Transcription Kit | ThermoFisher Scientific | 4368814 |
| Cells | | |
| VU147T | Hans Joenje, VU Medical Center, Netherlands | N/A |
| SCC4 | ATCC | CRL-1624 |
| SCC15 | ATCC | CRL-1623 |
| SCC90 | ATCC | CRL-3239 |
| SCC152 | ATCC | CRL-3240 |
| UMSCC47 (SCC47) | Millipore Sigma | SCC071 |
| OCTT2 | Devraj Basu, UPenn (Basu et al., 2010) | N/A |
| Primary gingival keratinocytes | ATCC | PCS-200-014 |
| Primers for qPCR | | |
| TaqMan Primers for T2R1 | ThermoFisher Scientific | Hs00251930_s1 |
| TaqMan Primers for T2R3 | ThermoFisher Scientific | Hs00249942_s1 |
| TaqMan Primers for T2R4 | ThermoFisher Scientific | Hs00249946_s1 |
| TaqMan Primers for T2R5 | ThermoFisher Scientific | Hs01549633_s1 |
| TaqMan Primers for T2R7 | ThermoFisher Scientific | Hs00256778_s1 |
| TaqMan Primers for T2R8 | ThermoFisher Scientific | Hs00256766_s1 |
| TaqMan Primers for T2R9 | ThermoFisher Scientific | Hs00256757_s1 |
| TaqMan Primers for T2R10 | ThermoFisher Scientific | Hs00256794_s1 |
| TaqMan Primers for T2R13 | ThermoFisher Scientific | Hs00256781_s1 |
| TaqMan Primers for T2R14 | ThermoFisher Scientific | Hs00256800_s1 |
| TaqMan Primers for T2R16 | ThermoFisher Scientific | Hs00249955_s1 |
| TaqMan Primers for T2R19 | ThermoFisher Scientific | Hs05000933_s1 |
| TaqMan Primers for T2R20 | ThermoFisher Scientific | Hs00604340_s1 |
| TaqMan Primers for T2R30 | ThermoFisher Scientific | Hs03054740_sH |
| TaqMan Primers for T2R31 | ThermoFisher Scientific | Hs00604313_sH |
| TaqMan Primers for T2R38 | ThermoFisher Scientific | Hs00604294_s1 |
| TaqMan Primers for T2R39 | ThermoFisher Scientific | Hs00603443_s1 |
| TaqMan Primers for T2R40 | ThermoFisher Scientific | Hs00602589_s1 |
| TaqMan Primers for T2R41 | ThermoFisher Scientific | Hs00603461_s1 |
| TaqMan Primers for T2R42 | ThermoFisher Scientific | Hs00704057_s1 |
| TaqMan Primers for T2R43 | ThermoFisher Scientific | Hs00853105_sH |
| TaqMan Primers for T2R45 | ThermoFisher Scientific | Hs00820227_s1 |
| TaqMan Primers for T2R46 | ThermoFisher Scientific | Hs00853124_s1 |
| TaqMan Primers for T2R50 | ThermoFisher Scientific | Hs00604351_s1 |
| TaqMan Primers for T2R60 | ThermoFisher Scientific | Hs00603474_s1 |
| TaqMan Primers for UBC | ThermoFisher Scientific | Hs01871556_s1 |
| Recombinant DNA | | |

| | | |
|---|---|---|
| nls-R-GECO | Addgene | 32462 |
| pECFP-DEVD-Venus | Addgene | 24537 |
| pECFP-DEVG-Venus | Addgene | 34538 |
| pcDNA3-FlipGFP-T2A-mCherry | Addgene | 124434 |
| pCX-SpiCee-NLS | Addgene | 140900 |
| pCX-SpiCee-NES | Addgene | 140901 |
| Software | | |
| MetaFluor | Molecular Devices | N/A |
| MetaMorph | Molecular Devices | N/A |
| QuantStudio 5 | Applied Biosystems, Inc | N/A |
| Prism v8 | GraphPad Software | N/A |
| ImageJ/FIJI | Open Source [1] | N/A |
| cBio Cancer Genomics Portal | cbioportal.org (Cerami et al., 2012; Gao et al., 2013) | N/A |

**Supplementary Table S2** Clinical data for HNSCC patients

| Subject | Age | Sex | Race | Primary Site | Subsite | p16 status | Pathologic stage |
|---|---|---|---|---|---|---|---|
| 1 | 69 | F | white | oral cavity | oral tongue | n/a | T1N0M0 |
| 2 | 62 | M | white | oral cavity | oral tongue | n/a | T2N0M0 |
| 3 | 72 | M | white | oral cavity | retromolar trigone | n/a | T2N0M0 |
| 4 | 65 | M | white | oropharynx | base of tongue | p16+ | T2N3M0 |
| 5 | 59 | M | white | oropharynx | base of tongue | p16+ | T3N0M0 |
| 6 | 52 | F | white | oropharynx | base of tongue | p16+ | T4N2M0 (recurrent) |
| 7 | 65 | M | white | oropharynx | tonsil | p16+ | T2N0M0 |
| 8 | 57 | M | white | oropharynx | tonsil | p16+ | T2N2M0 |
| 9 | 66 | M | white | oropharynx | parapharyngeal space[a] | p16- | T1N1M0 (recurrent) |
| 10 | 64 | M | white | oropharynx | tonsil | n/a | T2N2bM0 |

Clinical data for head and neck squamous cell carcinoma (HNSCC) patients included in quantitative PCR (qPCR) taste receptor expression analysis (Figure 2).Includes 3 patients with oral cavity cancer and 7 patients with oropharyngeal cancer. Pathologic staging based on the American Joint Committee on Cancer (AJCC) 8$^{th}$ edition TNM staging [2]. [a]Primary tumor was located in the tonsil with recurrence/metastasis to a parapharyngeal space lymph node.

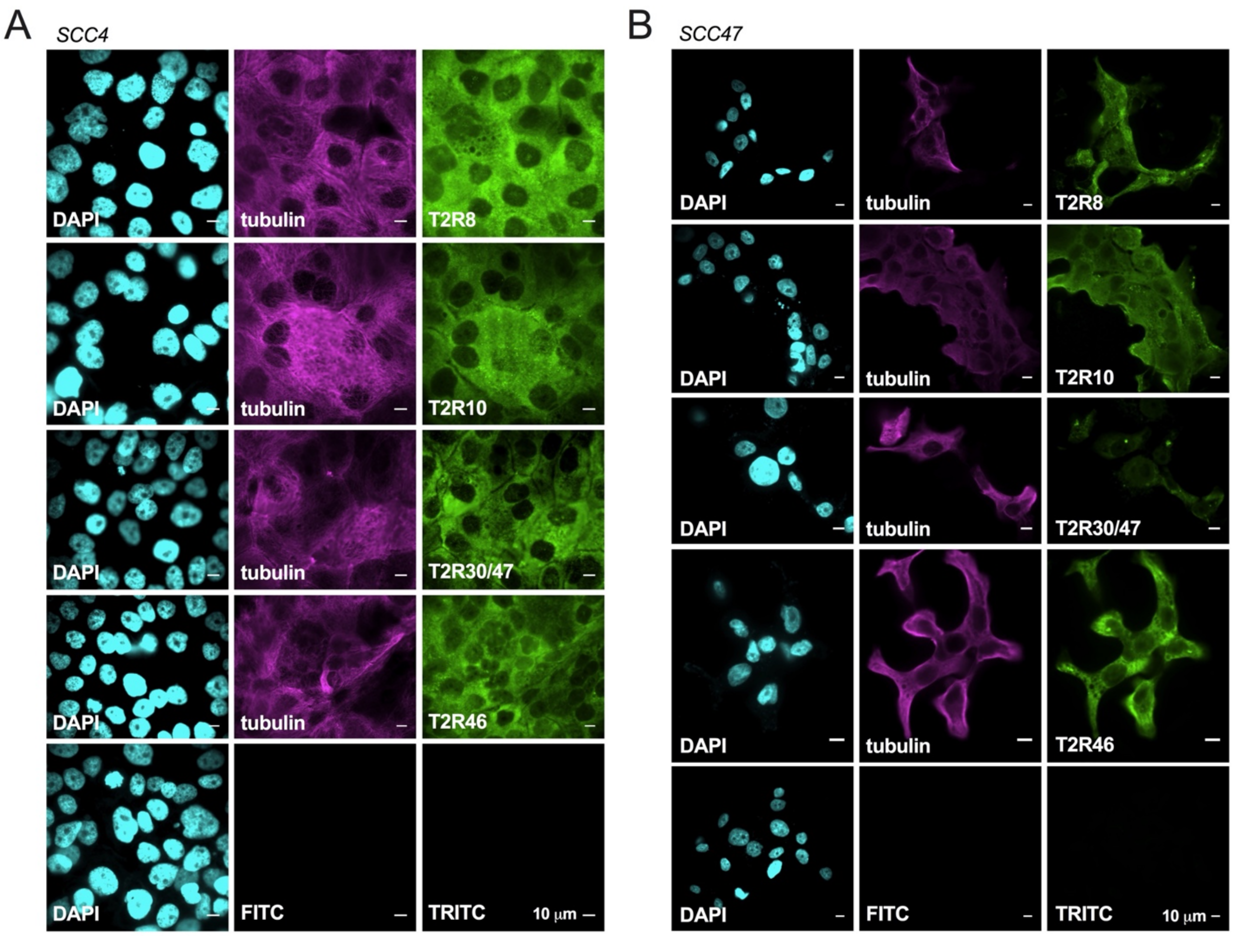


**Supplementary Fig. S1  T2Rs are expressed in head and neck squamous cell carcinoma (HNSCC) cell lines.** Fixed cultures of HNSCC cell lines SCC4 (A) and SCC47 (B) stained with antibodies targeting endogenous proteins demonstrate that T2Rs 8, 10, 30/47, and 46 localize to the plasma membrane. For all images, 1 representative image from 3 experiments were shown. Each antibody was compared to secondary only control at the same microscope settings.

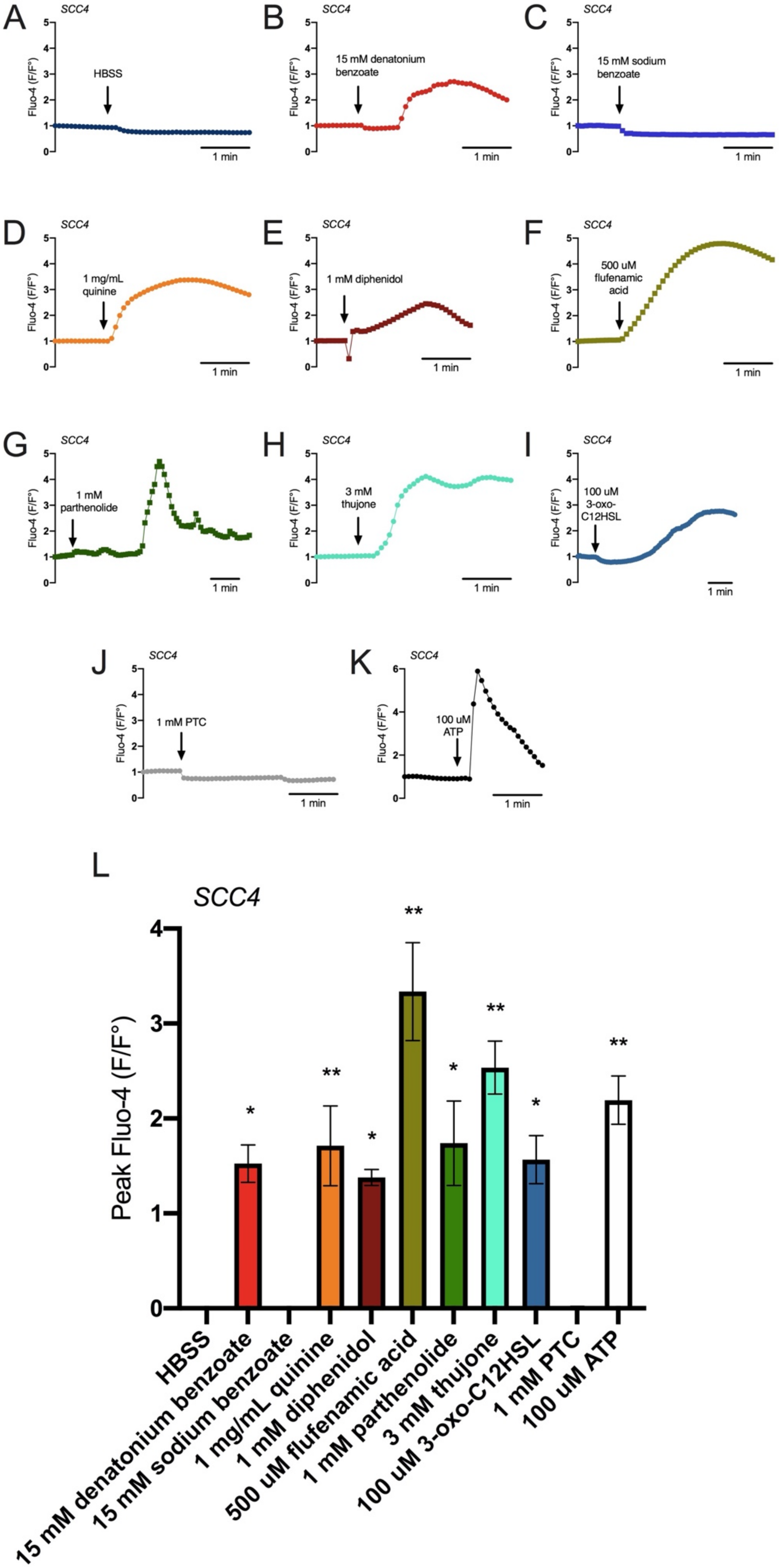
A
SCC4
Fluo-4 (F/F°)
HBSS
1 min
B
SCC4
15 mM denatonium benzoate
C
SCC4
15 mM sodium benzoate
D
SCC4
1 mg/mL quinine
E
SCC4
1 mM diphenidol
F
SCC4
500 uM flufenamic acid
G
SCC4
1 mM parthenolide
H
SCC4
3 mM thujone
I
SCC4
100 uM 3-oxo-C12HSL
J
SCC4
1 mM PTC
K
SCC4
100 uM ATP
L
SCC4
Peak Fluo-4 (F/F°)
HBSS
15 mM denatonium benzoate
15 mM sodium benzoate
1 mg/mL quinine
1 mM diphenidol
500 uM flufenamic acid
1 mM parthenolide
3 mM thujone
100 uM 3-oxo-C12HSL
1 mM PTC
100 uM ATP

**Supplementary Fig. S2 Bitter (T2R) agonists activate calcium responses in head and neck squamous cell carcinoma (HNSCC) cell line. A-K** HNSCC cell line SCC4 loaded with $Ca^{2+}$ binding dye, Fluo-4, was stimulated with T2R agonists and $Ca^{2+}$ was measured over time. Representative traces are shown after stimulation with Hank's Balanced Salt Solution (HBSS; control) (A), denatonium benzoate (B), sodium benzoate (C), quinine (D), diphenidol (E), flufenamic acid (F), parthenolide (G), thujone (H), N-3-oxo-dodecanoyl-L-homoserine lactone (3-oxo-C12HSL) (I), phenylthiocarbamide (PTC) (J), and purinergic receptor agonist adenosine triphosphate (ATP) (K). **L** Peak Fluo-4 $F/F_o$ was quantified and compared to HBSS (mean ± SEM; 3-7 experiments using separate cultures). Significance by 1-way ANOVA with Bonferroni post-test. Peak Fluo-4 $F/F_o$ was quantified after stimulation with Hank's Balanced Salt Solution (HBSS), denatonium benzoate, sodium benzoate, quinine, diphenidol, flufenamic acid, parthenolide, thujone, N-3-oxo-dodecanoyl-L-homoserine lactone (3-oxo-C12HSL), phenylthiocarbamide (PTC), and purinergic receptor agonist adenosine triphosphate (ATP) (mean ± SEM; 3-7 experiments using separate cultures). Significance by 1-way ANOVA with Bonferroni post-test comparing HBSS to each agonist. *$p<0.05$; **$p<0.01$.

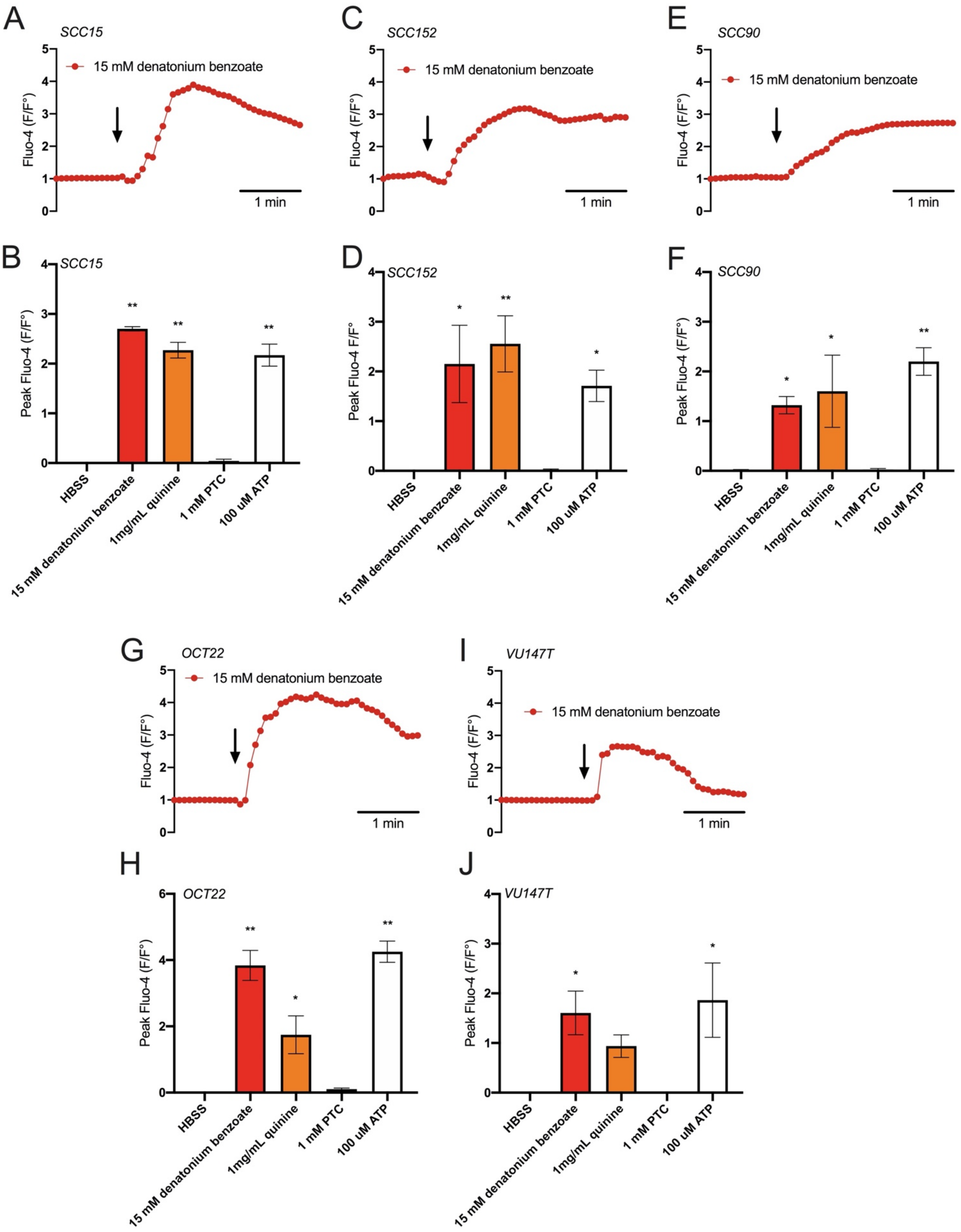
A
SCC15
15 mM denatonium benzoate
Fluo-4 (F/F°)
1 min
B
SCC15
Peak Fluo-4 (F/F°)
HBSS
15 mM denatonium benzoate
1mg/mL quinine
1 mM PTC
100 uM ATP
C
SCC152
15 mM denatonium benzoate
Fluo-4 (F/F°)
1 min
D
SCC152
Peak Fluo-4 F/F°
HBSS
15 mM denatonium benzoate
1mg/mL quinine
1 mM PTC
100 uM ATP
E
SCC90
15 mM denatonium benzoate
Fluo-4 (F/F°)
1 min
F
SCC90
Peak Fluo-4 (F/F°)
HBSS
15 mM denatonium benzoate
1mg/mL quinine
1 mM PTC
100 uM ATP
G
OCT22
15 mM denatonium benzoate
Fluo-4 (F/F°)
1 min
H
OCT22
Peak Fluo-4 (F/F°)
HBSS
15 mM denatonium benzoate
1mg/mL quinine
1 mM PTC
100 uM ATP
I
VU147T
15 mM denatonium benzoate
Fluo-4 (F/F°)
1 min
J
VU147T
Peak Fluo-4 (F/F°)
HBSS
15 mM denatonium benzoate
1mg/mL quinine
1 mM PTC
100 uM ATP

**Supplementary Fig. S3 Bitter (T2R) agonists activate calcium responses in head and neck squamous cell carcinoma (HNSCC).** HNSCC cell lines loaded with Fluo-4 were stimulated with denatonium benzoate (agonist for T2R4, 8, 10, 13, 39, 43, 46, and 47), quinine (agonist for T2R4, 7, 10, 14, 39, 40, 43, 44, and 46), phenylthiocarbamide (PTC; agonist for T2R38), and adenosine triphosphate (ATP; agonist for purinergic receptors) and calcium responses were measured over time. Representative traces after stimulation with denatonium benzoate from single cultures of SCC15 (A), SCC152 (C), SCC90 (E), OCT22 (G), and VU147T (I). Peak Fluo-4 $F/F_o$ was quantified for each cell line (SCC15 (B), SCC152 (D), SCC90 (F), OCT22 (H), and VU147T (J)) after stimulation with Hank's Balanced Salt Solution (HBSS), denatonium benzoate, quinine, PTC, and ATP (mean ± SEM; 3-7 experiments using separate cultures for each cell line). Significance by 1-way ANOVA with Bonferroni post-test comparing HBSS to each agonist. *$p<0.05$; **$p<0.01$.

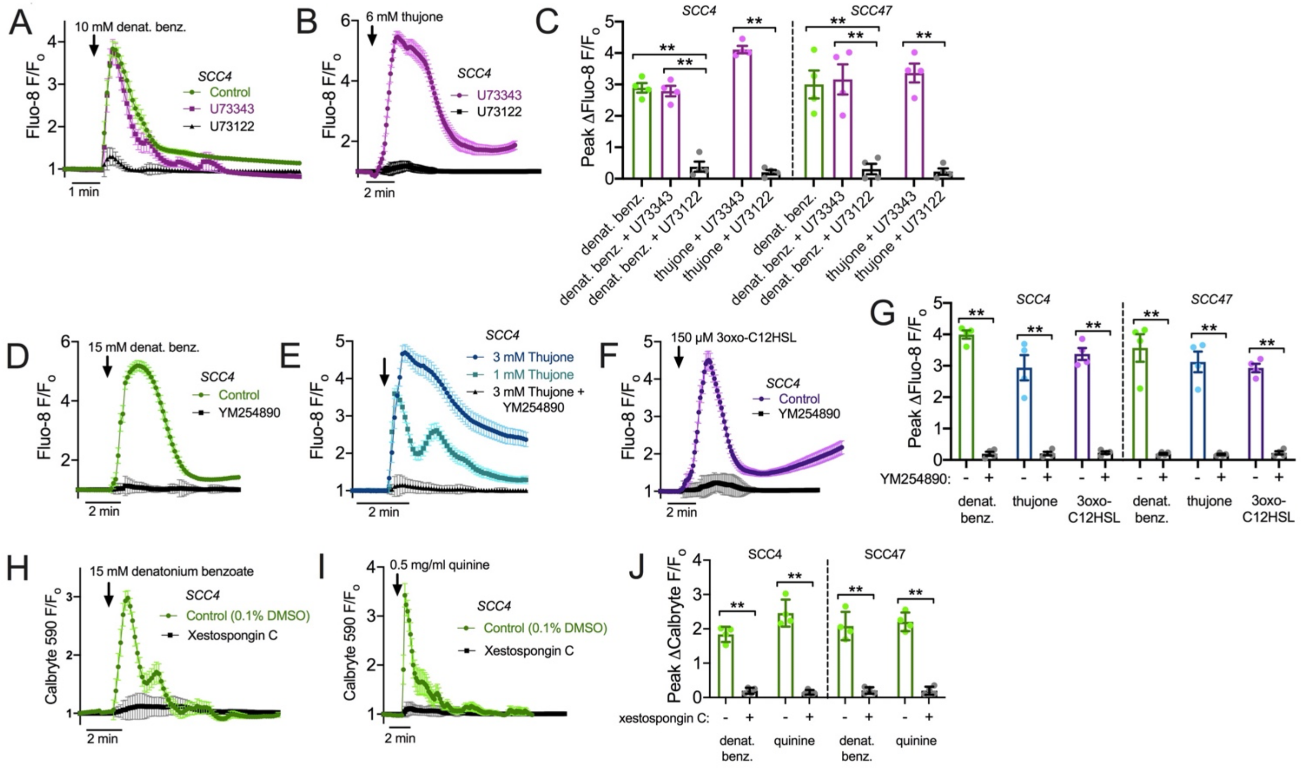


**Supplementary Fig. S4 Pharmacology of the $Ca^{2+}{}_i$ response. A-C** SCC4 and SCC47 cells were loaded with Fluo-8 for 45 min and imaged as described in the text. Cells were loaded in the presence of 0.1% DMSO (vehicle control), 10 µM phospholipase C inhibitor U73122, or 10 µM inactive analogue U73343. Cells were stimulated with denatonium benzoate or thujone as indicated in the continued presence or absence of inhibitor. Representative traces are shown from SCC4 for denatonium (A) and thujone (B). Bar graph ***(C)*** shows inhibition of $Ca^{2+}{}_i$ responses by U73122 but not U73343. **D-G** SCC4 and SCC47 cells were loaded with Fluo-8 for 45 min and imaged as described. After loading, cells were incubated with 0.1% DMSO (vehicle control) or 10 µM YM254890, a heterotrimeric G protein inhibitor. Cells were then stimulated with denatonium, thujone, or 3-oxo-C12HSL. Representative traces from SCC4 shown in D, E, and F, respectively. Bar graph (G) shows inhibition of $Ca^{2+}{}_i$ with all agonists in the presence of YM254890. **H-J** SCC4 and SCC47 cells were loaded with calcium indicator Calbryte 590 for 45 min and imaged as described in the text. Cells were pre-incubated for 10 min with 0.1% DMSO (vehicle control) or 10 μM xestospongin C, and $IP_3R$ inhibitor. H and I show representative traces from SCC4 cells stimulated with denatonium or quinine, respectively. Bar graph (J) shows reduced $Ca^{2+}_i$ response with xestospongin C. All traces shown are representative experiments showing mean ± SEM of 20-40 cells from a single field of view. Data points in bar graphs are independent experiments (n = 3-6). Significance determined by one-way ANOVA with Bonferonni posttest; $^{**}p<0.01$.

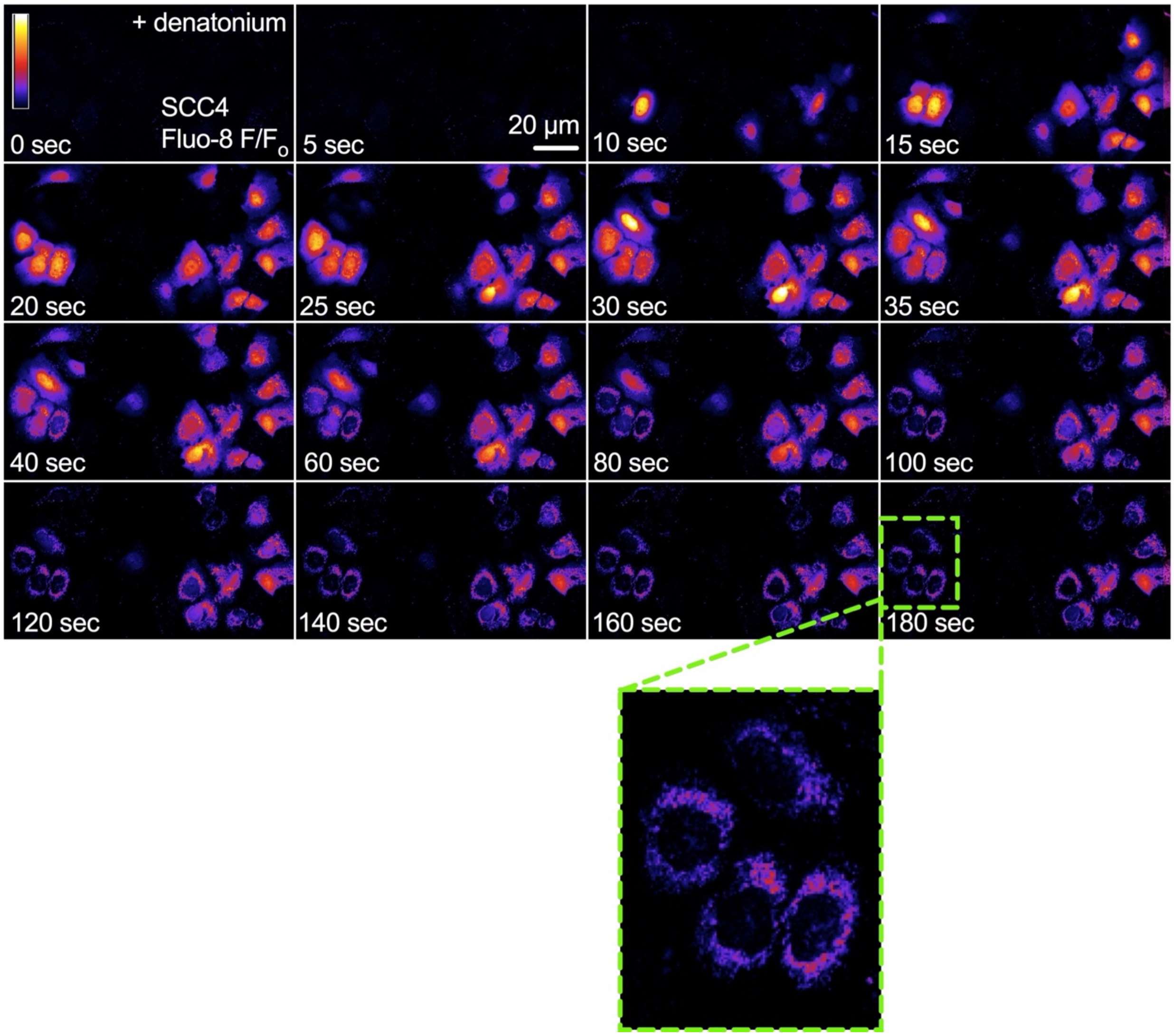


**Supplementary Fig. S5 Nuclear $Ca^{2+}$ responses appeared to propagate to mitochondria in Fluo-8-loaded cells.** Representative F/F$_o$ images of a time course of denatonium stimulation in SCC4 cells loaded with Fluo-8. The initial nuclear $Ca^{2+}$ increase was followed by a lower-level but more sustained increase in $Ca^{2+}$ in a perinuclear pattern reminiscent of mitochondria. Green outlined box at bottom shows enlargement of outlined region from 180 sec time point. These observations suggested that bitter agonist-activated nuclear $Ca^{2+}$ signals might influence mitochondrial function.

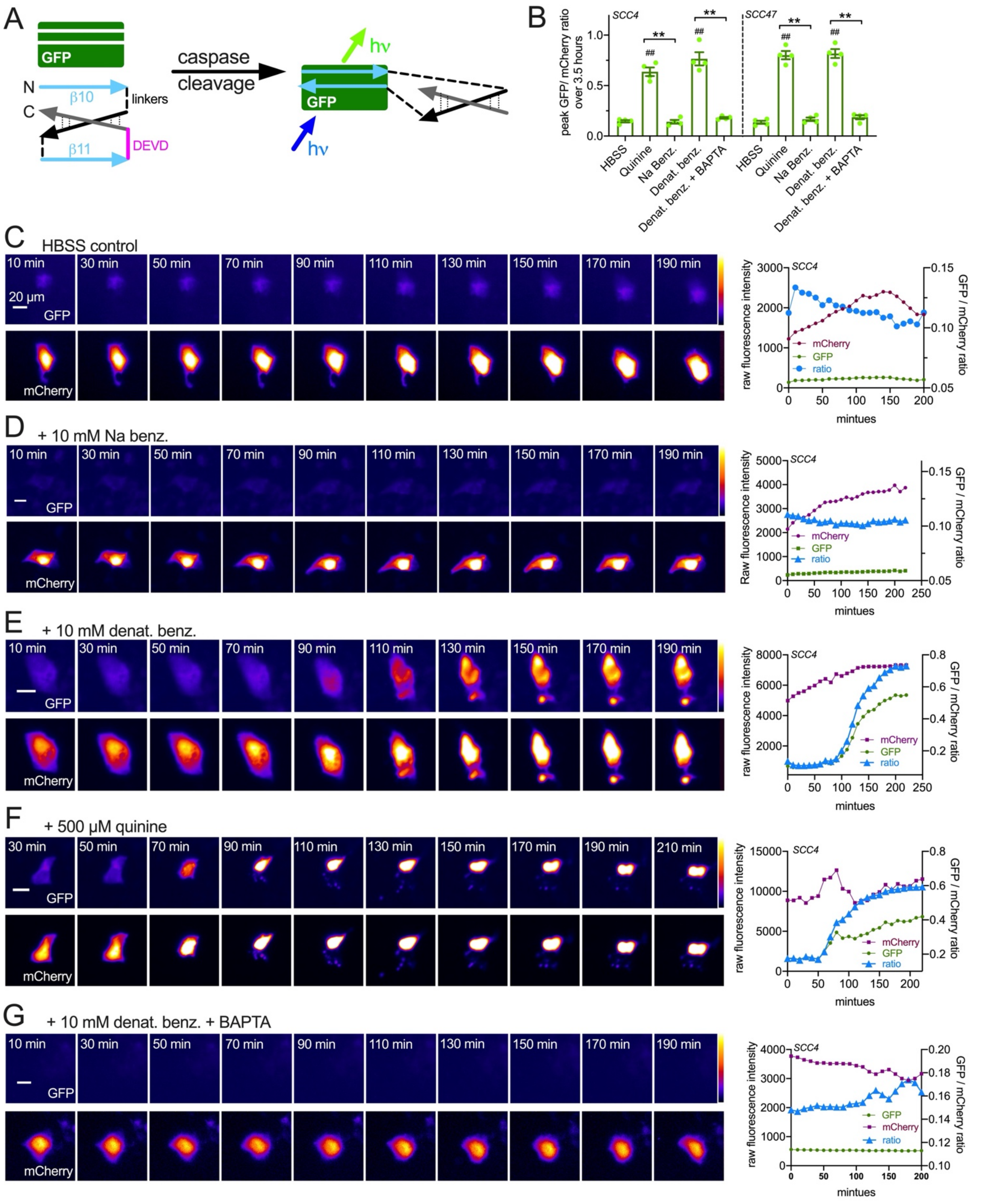
A
GFP
N
C
β10
β11
linkers
DEVD
caspase
cleavage
hν
GFP
hν
B
SCC4
SCC47
peak GFP/ mCherry ratio over 3.5 hours
HBSS
Quinine
Na Benz.
Denat. benz.
Denat. benz. + BAPTA
C HBSS control
20 µm
GFP
mCherry
raw fluorescence intensity
GFP / mCherry ratio
mintues
D + 10 mM Na benz.
E + 10 mM denat. benz.
F + 500 µM quinine
G + 10 mM denat. benz. + BAPTA

**Supplementary Fig. S6 Confirmation of caspase activation by Flip-GFP and dependence of denatonium-induced caspase activation on $Ca^{2+}$ signaling. A** Diagram of the Flip-GFP assay [3]. GFP β10 and β11 strands are anti-parallel within the GFP β barrel. The Flip construct contains parallel β10 and β11 joined by a linker. Cleavage of the DEVD sequence in the construct permits flipping of β11, allowing β10 and β11 to fit into the GFP barrel and complete the GFP. Thus, caspase activity will increase GFP fluorescence. Soluble mCherry is expressed as a transfection control. **B** Cells were imaged with a 10x objective and GFP and mCherry fluorescence was measured in single cells over 3-4 hours (representative experiments below). Background-subtracted fluorescence ratios are shown after 3.5 hours (210 min) stimulation as indicated. BAPTA-loaded cells were pre-incubated with 10 μM of global calcium chelator BAPTA-AM for 1 hour. Cells for all other conditions were similarly incubated in the absence of BAPTA). Each data point represents one cells from an independent experiment (n=3-5). Bar graph is mean ± SEM. Significance by one way ANOVA with Bonferonni posttest; ## $p<0.01$ vs HBSS control; ** $p<0.01$ vs bracketed bars. **C-G** Representative intensity pseudocolored images of GFP and mCherry fluorescence (left) as well as traces showing fluorescence changes in SCC4 cells during incubation with HBSS (control; *C*), sodium benzoate (*D*), denatonium benzoate (*E*), quinine (*F*), or denatonium + BAPTA (*G*). Background subtraction and quantification of cell fluorescence was carried out using ImageJ.

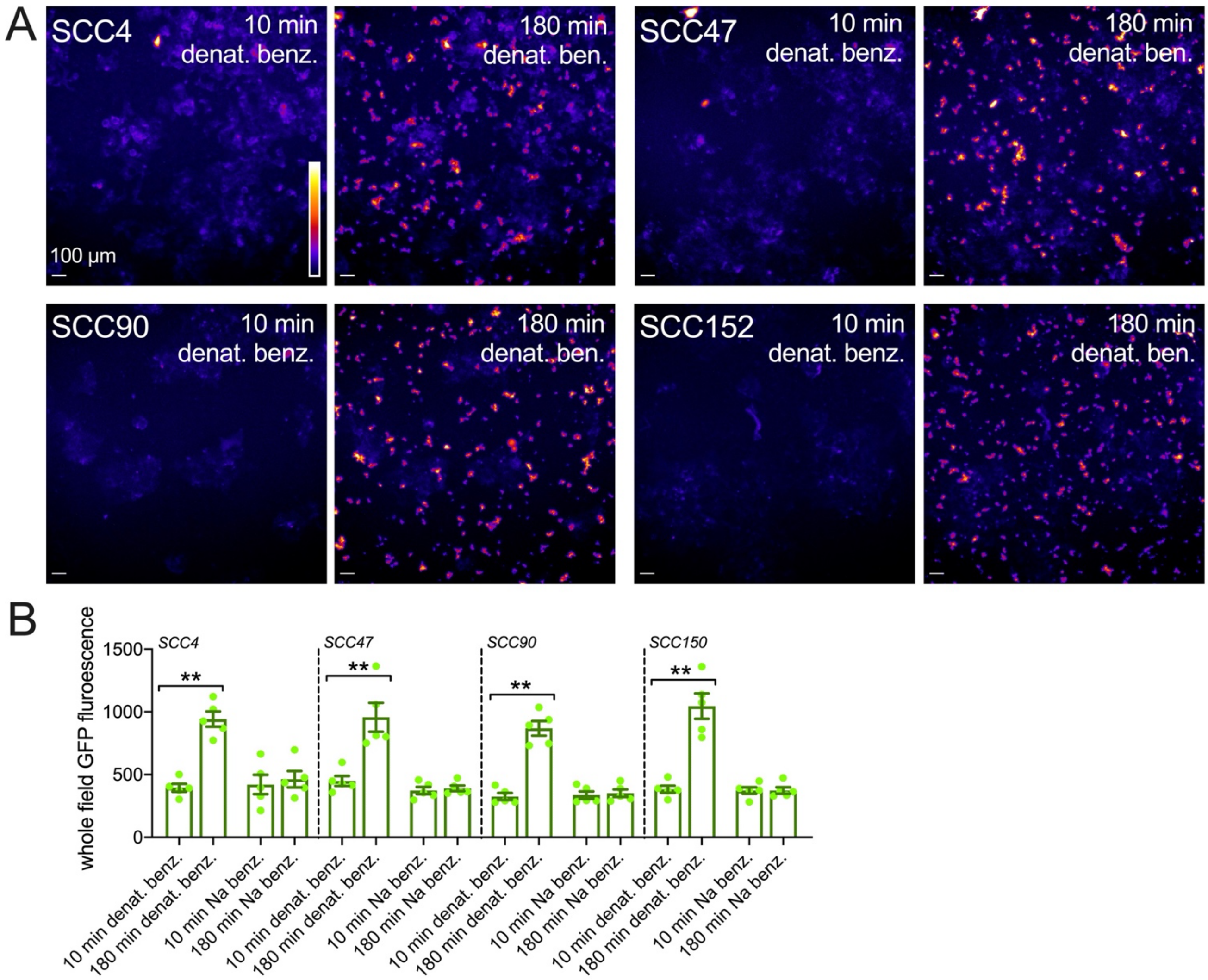


**Supplementary Fig. S7 Flip-GFP measurement of caspase activation in response to denatonium benzoate but not sodium benzoate in SCC4, SCC47, SCC90, and SCC152.** Cells were imaged using 4x objective and GFP fluorescence was estimated from whole field fluorescence intensity. **A** Representative intensity pseudocolored images of GFP fluorescence at 10 min and 180 min after denatonium benzoate stimulation in four different cell lines as indicated. **B** Bar graph of quantified data from experiments in *(A)* where cells were stimulated with 10 mM denatonium benzoate or sodium benzoate. Each data point represents one cells from an independent experiment (n=3-5). Bar graph is mean ± SEM. Significance by one way ANOVA with Bonferonni posttest; ** p<0.01 vs bracketed bars. Background subtraction and quantification of cell fluorescence was carried out using ImageJ.

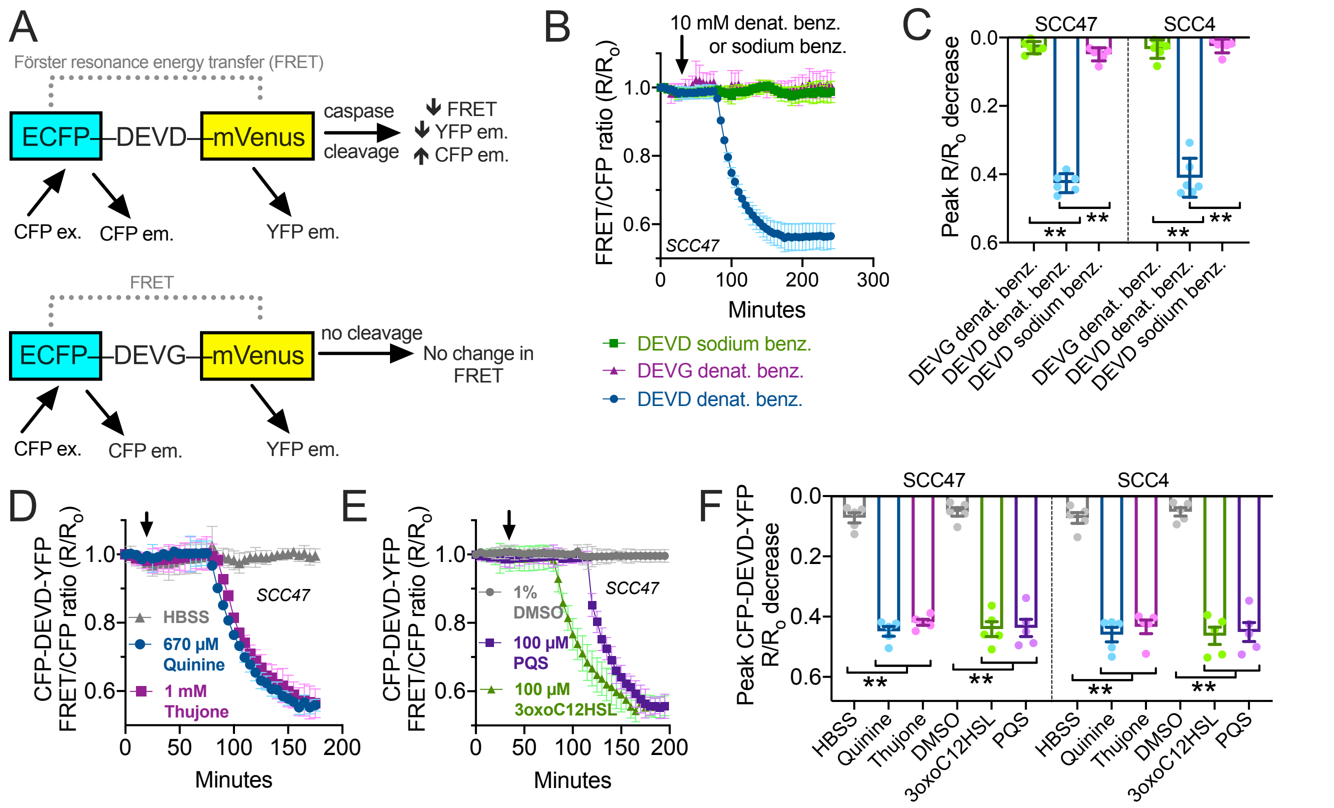


**Supplementary Fig. S8 Confirmation of bitter agonist-induced caspase activation by ratiometric caspase biosensor.** **A** Schematic of the biosensor (top) showing enhanced CFP (ECFP) connected to YFP variant mVenus by a DEVD linker. Caspase cleavage allows the ECFP and mVenus to diffuse apart and reduces FRET signal (YFP emission with CFP excitation). A control biosensor (bottom) was used with the DEVD replaced with a non-cleavable DEVG. **B** Representative traces (mean ± SEM of 5-9 transfected cells) from SCC47 cells showing decreased FRET after ~60 min denatonium benzoate stimulation with DEVD biosensor but not uncleavable control DEVG biosensor. Sodium benzoate had no effect. **C** Bar graph showing peak FRET decrease in SCC47 and SCC4 cells from independent experiments as in *B*. **D-E** Representative traces of DEVD biosensor FRET ratio in SCC47 cells stimulated with quinine (D), thujone (D), Pseudomonas quinolone signal (PQS; (E)), 3-oxo-C12HSL (E), or 1% DMSO (vehicle control for 3-oxo-C12HSL and PQS; (E)). **F** Bar graph showing peak FRET decrease in SCC47 and SCC4 cells from independent experiments as in (D-E). Data points in bar graphs are independent experiments (n = 3-6). Significance determined by one-way ANOVA with Bonferonni posttest; ***p*<0.01. Together, these data support activation of caspases by multiple bitter agonists, including bacterial 3-oxo-C12HSL and PQS.

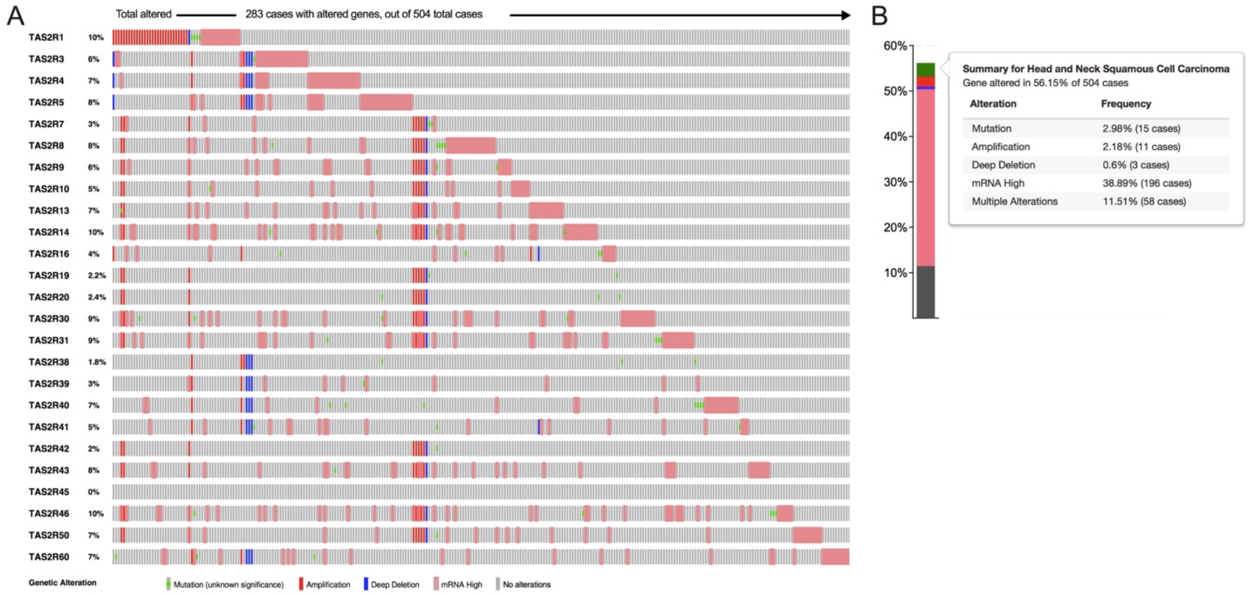


**Supplementary Fig. S9 Bitter taste receptor (*TAS2R*) genomic and expression alterations are prevalent in head and neck squamous cell carcinoma (HNSCC).** *TAS2R* genetic and expression alterations were analyzed for 504 cases of HNSCC using The Cancer Genome Atlas (TCGA) [4, 5]. A total of 283 out of 504 cases (56.15%) included some *TAS2R* genetic or expression alteration. **A** OncoPrint of *TAS2R* genomic and expression alterations in HNSCC. Genes are listed in rows with percentage of altered cases; individual cases are listed in columns. **B** Bar graph demonstrating alteration types and frequencies in HNSCC. Multiple alterations were present in 11.51% of cases.

### Supplementary References


1. Schindelin J, Arganda-Carreras I, Frise E, Kaynig V, Longair M, Pietzsch T*, et al.* Fiji: an open-source platform for biological-image analysis. *Nat Methods* 2012, **9**(7)**:** 676-682.

2. Amin MB, Greene FL, Edge SB, Compton CC, Gershenwald JE, Brookland RK*, et al.* The Eighth Edition AJCC Cancer Staging Manual: Continuing to build a bridge from a population-based to a more "personalized" approach to cancer staging. *CA Cancer J Clin* 2017, **67**(2)**:** 93-99.

3. Zhang Q, Schepis A, Huang H, Yang J, Ma W, Torra J*, et al.* Designing a Green Fluorogenic Protease Reporter by Flipping a Beta Strand of GFP for Imaging Apoptosis in Animals. *J Am Chem Soc* 2019, **141**(11)**:** 4526-4530.

4. Cerami E, Gao J, Dogrusoz U, Gross BE, Sumer SO, Aksoy BA*, et al.* The cBio cancer genomics portal: an open platform for exploring multidimensional cancer genomics data. *Cancer discovery* 2012, **2**(5)**:** 401-404.

5. Gao J, Aksoy BA, Dogrusoz U, Dresdner G, Gross B, Sumer SO*, et al.* Integrative analysis of complex cancer genomics and clinical profiles using the cBioPortal. *Sci Signal* 2013, **6**(269)**:** pl1.
